# Cross-regulatory sRNA networks shape adaptive stress responses in *Staphylococcus aureus*: RsaC and RsaOI regulate galactose metabolism and cell wall homeostasis

**DOI:** 10.64898/2026.10.30.763024

**Authors:** Hugo Robin, Maëliss Germain, Chloé Silard, Stéphane Dreano, Yoann Augagneur, Svetlana Chabelskaya, Astrid Rouillon

**Affiliations:** Université de Rennes, BRM, Bacterial RNAs and Medicine, UMR_S 1230, Rennes, France; Université de Rennes, CNRS, IGDR, Institute of Genetics and Development of Rennes, BIOSIT, Rennes, France

**Keywords:** *Staphylococcus aureus*, sRNA, regulatory network, cell wall stress, SpA

## Abstract

*Staphylococcus aureus* is a major human pathogen whose adaptability relies on network of regulators, including two-component systems, transcription factors and small regulatory RNAs. Yet most of these regulators are dispensable for growth, suggesting cooperation or redundancy between them. To explore this hypothesis, we performed a yeast one-hybrid screen of 43 TFs against the promoters of seven sRNAs, which identified 27 novel TF-sRNA promoter interactions. We then focused on three sRNAs. We showed that *srn_9342* is repressed by VraR and MntR, which also control *rsaOI* and *rsaC*, respectively. We found that *rsaC*, like *rsaOI*, is induced by acidic pH and vancomycin exposure, whereas *srn_9342* expression remained unchanged. We also demonstrated *in vitro* that Srn_9342 interacts with both RsaOI and RsaC. Its role toward these two sRNAs remains unknown, but this interaction raises the possibility that it modulates their availability or activity. RsaOI and RsaC both act on the tagatose pathway for galactose utilization, through different genes: *lacD* for RsaOI, as previously reported, and *lacC* for RsaC, newly identified here. We showed that both sRNAs are required for surface charge remodeling and for growth under subinhibitory vancomycin. They also modulate the virulence factors PSMα and protein A. Both sRNAs reduce PSMα production, whereas protein A is decreased by RsaOI, likely through Rot, and increased by RsaC through an unknown, probably indirect mechanism. Together, our results suggest that a single cell wall stress can engage several regulators that could cooperate on shared targets or pathways, which may explain why individual regulators are dispensable.

## Introduction

*Staphylococcus aureus* is a Gram-positive commensal bacterium, carried asymptomatically by approximately 30% of the human population on their skin and in their nostrils, which constitutes the main reservoir for subsequent infections (1). However, under conducive conditions, it can behave as a major opportunistic pathogen responsible for a spectrum of diseases, ranging from superficial skin lesions to life-threatening infections such as endocarditis, pneumonia, and toxic shock syndrome (2).

The success of *S. aureus* as a pathogen relies in large part on its remarkable adaptability, enabling it to survive in diverse environments, evade the immune system, and withstand antibiotic treatment. This adaptability depends on its ability to sense and respond to environmental changes by modulating gene expression, using multiple regulators, including two-component systems (TCSs), transcription factors (TFs) and small regulatory RNAs (sRNAs).

*S. aureus* encodes 16 conserved TCS that govern physiological processes as diverse as virulence, antibiotic tolerance, cell wall metabolism, respiration, and nutrient sensing (3). These regulatory circuits are autonomous modules that sense and respond to specific environmental cues, although some level of cross-regulation between non-cognate sensor-response regulator pairs has been observed *in vivo* (4). Specific TCSs have been implicated in various critical functions, including cell wall metabolism, autolysis, biofilm development, and cell death (WalKR, ArlSR, and LytSR), to bacterial respiration, fermentation, and nitrate metabolism (SrrBA, AirSR, and NreBC), as well as nutrient sensing and metabolism (HssRS, KdpDE, and PhoRP) (5, 6). Additionally, TCSs play significant roles in virulence (AgrCA and SaeSR) (7–10) and antibiotic resistance and cell wall damage (VraSR, GraXSR, and BraSR) (11, 12). Among them, WalKR is the only essential system: the fifteen others can be deleted concomitantly from the same strain without compromising cellular viability (5).

In addition to TCS, which regulate transcription as paired entities, TF act independently to modulate gene expression. In 2013, a study of TF-encoding genes across *S. aureus* strains identified 120 TFs distributed in 34 regulator families, nearly half of which had no assigned function at the time (13). Among these families, the MarR family stands out as the largest, with 18 members including 10 proteins of the Sar subfamily (SarA, SarR, SarS, SarT, SarU, SarV, SarX, SarZ, MgrA, and Rot) (14).

Despite this diversity, these regulators converge on common targets, and it is their combined, cooperative, or antagonistic interactions that ultimately shape gene expression. A prime example of this regulatory interplay is provided by the production of phenol-soluble modulins (PSMs), which are key virulence factors in *Staphylococcus aureus*. In this system, the two-component system AgrC/A enhances the expression of *psmα* and *psmϐ* (15), whereas the transcription factors MgrA and SarA act as respectively repressor or activator of *psm* production (16, 17). This antagonistic regulatory interplay is not unique, the gene *spa*, encoding protein A, provides a second example, being activated by SarS and Rot and repressed by SarA and MgrA (18–20). Regulators also act on one another: SarA represses *mgrA* transcription (21) and autoregulates its own expression (22). Such overlapping and frequently antagonistic inputs contribute to the fine-tuned control of bacterial physiology but also present challenges in deciphering individual regulator functions.

In addition to these transcription factors, small RNAs (sRNAs) represent a supplementary layer of post-transcriptional regulation. The repertoire of sRNAs in *S. aureus* varies depending on the identification criteria employed. In 2015, a comprehensive review of existing data compiled a list of 575 potential sRNA sequences (23). When only sRNAs with at least one directly identified target are considered, this number falls to approximately 15 sRNAs (24). Liu and colleagues defined *bona fide* sRNAs as those that (i) likely act in trans and (ii) are not expressed from the opposite strand of a coding gene. This classification allowed them to identify 50 *bona fide* sRNAs (25). Among these, SprD and RsaD have well-characterized targets, repressing Sbi and AlsS expression, respectively, and playing roles in bacterial virulence and metabolism (26, 27). Although sRNAs regulate targets involved in bacterial virulence or metabolism, none has been found to be essential for bacterial growth.

These three classes of regulators do not act independently but are tightly interconnected within regulatory networks (28). In particular, sRNAs are both targets and effectors of TFs. For instance, SarA, a major TF in *Staphylococcus aureus*, directly represses the expression of several sRNAs, including *sprC*, *srn_9340* and *rsaD*, the latter being also controlled by CodY (21, 27). This dual control appears to be growth-phase dependent: CodY-mediated repression of *rsaD* likely occurs at low cell density, whereas SarA-dependent repression takes place later (21). Conversely, sRNAs can feed back on these protein regulators, RsaA and RsaC downregulate MgrA and SarA expression respectively (29, 30), thereby closing regulatory loops. The extent of this interconnection becomes clearer when the size of the regulatory repertoire is compared with that of the genome. In *S. aureus*, approximately 120 TFs have been identified, either experimentally or by sequence homology, representing less than 5% of the genes (13). With ∼533 operons (31) covering 60% of the genome (32), the remaining ∼40 % of genes being monocistronic, the genome is estimated to contain ∼1,700 promoters, so that each TF would control, on average, approximately 14 promoters. The inactivation of a single regulator would therefore be expected to cause major perturbations of gene expression and strong phenotypes. Yet the vast majority of these regulators are dispensable for bacterial growth, including 15 of the 16 TCSs and all characterized sRNAs.

These observations raise the possibility that regulators compensate for one another by acting on shared targets. sRNAs are well suited to test this hypothesis, as some of them integrate inputs from several regulators, as illustrated by *rsaD*, which is repressed by both SarA and CodY (21, 27). Whether such co-regulation is widespread, and whether it could explain functional redundancy, remains largely unexplored. Here, we therefore aimed to identify the protein regulators binding sRNA promoters and to determine whether shared regulators could account for the dispensability of each of them individually.

Using a heterologous one-hybrid screen in *Saccharomyces cerevisiae* (Y1H), we tested 43 regulators (TFs and TCS response regulators) against the promoters of seven *S. aureus* sRNAs. The screen revealed 29 interactions, 27 of them previously undescribed, with some regulators binding up to three promoters, thus identifying sRNAs that share common regulators. After validating these interactions in *S. aureus*, we focused on Srn_9342, RsaOI and RsaC. *srn_9342* was repressed by VraR and MntR, two regulators already known to control *rsaOI* and *rsaC*, respectively (33, 34), and its interactions with both sRNAs, previously suggested by MAPS experiments (35), were confirmed *in vitro*. *rsaOI* is known to be induced by acidic pH and vancomycin (33), we show that *rsaC* is also induced under both conditions, whereas *srn_9342* transcription remains unchanged. We further show that RsaC targets the *lacC* mRNA and impairs galactose utilization, as previously reported for RsaOI through repression of *lacD* (33), suggesting that these two sRNAs jointly regulate carbon metabolism by targeting different genes of the same operon. We also found, most likely through indirect mechanisms, that *rsaOI* overexpression decreased and *rsaC* overexpression increased protein A expression, whereas overexpression of either sRNA repressed *psmα* expression, at both the transcript and protein levels. Together, these results suggest that sRNAs controlled by different regulators converge on shared pathways and/ or targets (galactose utilization, protein A, *psmα*), providing a possible mechanism for functional redundancy and cooperation between regulators.

Together, these results support a model in which sRNAs integrate inputs from different regulators and converge on shared pathways and targets, providing a possible mechanism for functional redundancy and cooperation between regulators.

## Results

### Decoding sRNA gene regulation: Yeast One-Hybrid screening in *Saccharomyces cerevisiae* discovers candidate transcription factors

The yeast one-hybrid assay allows the evaluation of a protein’s ability to recognize and bind to a specific DNA sequence, referred to as bait (36, 37). In this method, the protein of interest (i.e. TF) is fused to a yeast transcriptional activation domain (here, Gal4_AD_) and is referred to as the TF-prey. When the TF-prey binds the bait sequence, Gal4_AD_ recruits the transcription machinery and drives expression of a reporter gene (*HIS3* in this study), allowing yeast growth on histidine-free medium (Figure 1-A). The assay therefore identifies the intrinsic DNA-binding capacity of a TF, but do not reveal its positive or negative regulatory effect. In this study, we adapted Y1H to identify transcriptional regulators for several sRNA genes in *S. aureus*.

**Figure 1.**
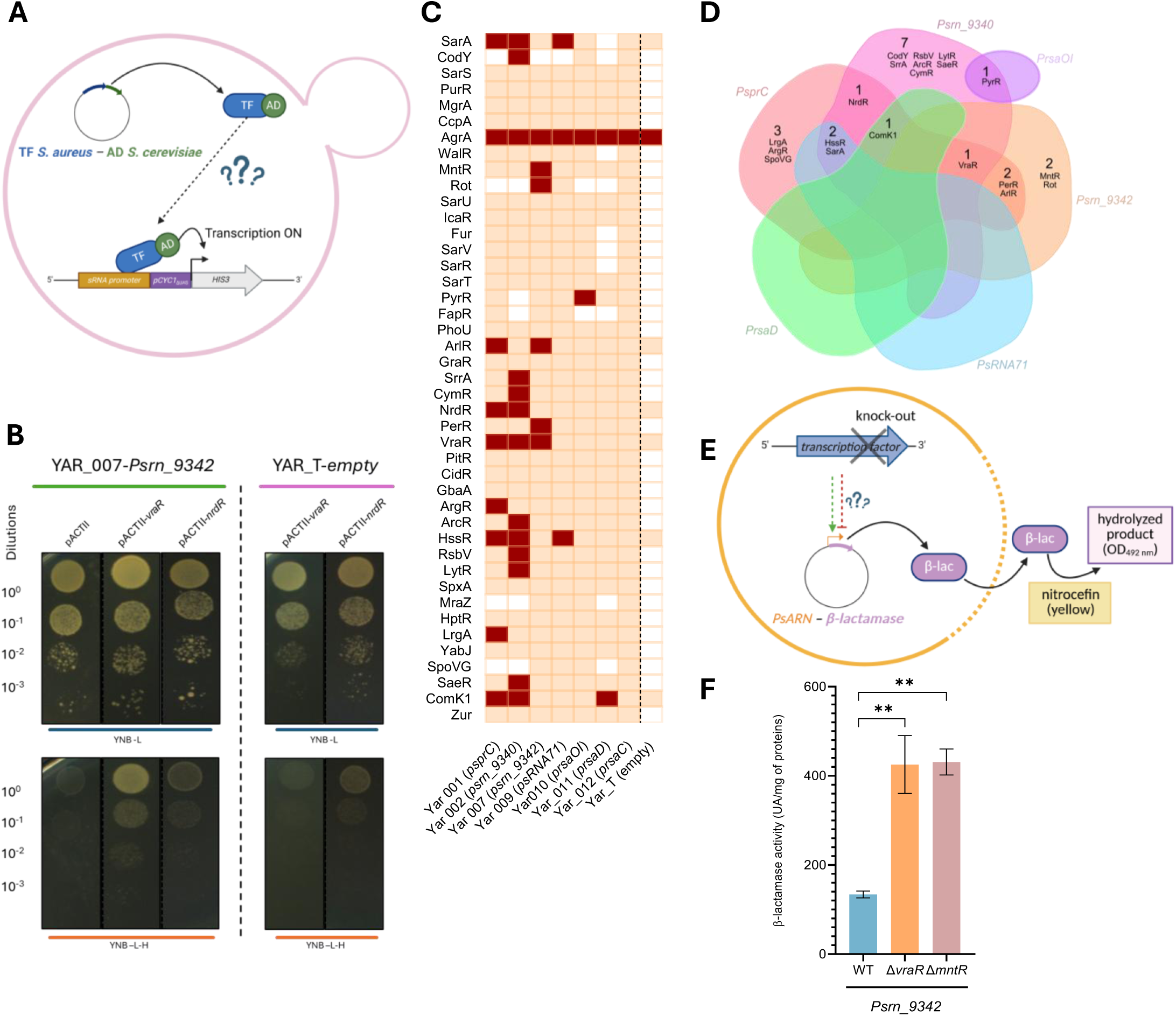
Y1H screening revealed promoters-TFs binding occurring in *S. aureus*. (**A**) Schematic representation of the Y1H screening assay, promoter–TF interactions are revealed by histidine prototrophy. (**B**) Representative spot assay showing promoter-TF interaction on selective medium. (**C**) Heatmap representing the results of the Y1H screening for all tested sRNA promoter–TF pairs (red: TF–sRNA promoter interaction; orange: no interaction; white: no transformant). (**D**) Venn diagram of positive TF hits. (**E**) Schematic of the β-lactamase reporter assay used for functional validation in a TF deletion context in *S. aureus,* transcriptional repressors and activators are detected by an increase or decrease in OD_492 nm_, respectively, compared to WT. (**F**) Validation of two candidates TFs, VraR and MntR, on the *srn_9342* promoter using the β-lactamase reporter assay. Cells were grown for 5h in TSB medium prior to the assay. Statistical analysis was conducted using Student’s *t*-test. Error bars represent the average of three independent experiments. Statistical significance is indicated by bars and asterisks as follows: \*\**P*< .01.

Of the 120 transcriptional regulators annotated in *S. aureus* (13), we prioritized transcription factors (TFs) that possessed experimentally validated regulatory roles on at least one target gene or signaling pathway. By integrating these pre-selected transcriptional regulators candidates into our Y1H analysis, we aimed to identify novel *srna* promoter interactions and contextualize them within well-characterized transcriptional networks. Therefore, 43 TFs from *S. aureus*, including 10 response regulators of TCSs, were selected for cloning into the pACTII vector in fusion with the Gal4_AD_ of yeast creating 43 TF-prey plasmids (Table S1). Regarding the *srna* promoters, our choice was guided by those previously studied in our laboratory SprC, Srn_9340, Srn_9342, RsaOI, RsaD (27, 33, 35, 38) and two others sRNA71 and RsaC (30, 39). The promoter regions of these sRNAs were individually cloned upstream of the *HIS3* reporter gene within the pCyc1_ΔUAS_ yeast promoter, creating the seven yeast DNA-bait reporter strains (referred as YAR-strain in Table S1).

Following yeast transformation and clone selection, the growth on media with and without histidine was assessed to identify the TFs capable of interacting with sRNA promoter sequences. To evaluate the specificity of the yeast one-hybrid (Y1H) assay, we first transformed each YAR strain with the empty pACTII vector and verified, for each sRNA promoter tested, that Gal4_AD_ alone did not produce a positive signal. Next, when a given TF prey produced a positive signal in at least two YAR strains, it was transformed into the YAR_T-empty strain. This control strain carries the pCYC1_ΔUAS_ yeast promoter without any cloned sRNA promoter, and therefore serves as a negative control to detect non-specific activation of the reporter by *S. aureus* TFs. These controls allowed us to distinguish bait-dependent interactions from non-specific reporter activation.

We thus obtained and tested 279 individual combinations out of the 301 envisioned. Some TF-preys proved difficult to transform, such as CodY, Rot, FapR, MraZ, and SpoVG, though the underlying reasons for these difficulties remain unclear. For each combination, we compared growth with and without histidine. For example, in the Yar_007 strain, neither pACTII (empty vector) nor pACTII_NrdR allowed growth without histidine, whereas pACTII_VraR did. As the Yar_T-empty strain (lacking the bait sequence) carrying either Gal4_AD_-VraR or Gal4_AD_-NrdR was unable to grow without histidine, it confirms that the interaction is specific to the bait sequence. We therefore conclude that VraR binds the promoter of *srn_9342*, whereas NrdR does not (Figure 1-B). Among the combinations tested, 36 strains were capable of growing without histidine, involving 20 distinct TFs (Figure 1-C). Out of these 20 positives TFs, AgrA was eliminated from further study as it was found to activate the transcription of the reporter gene in the Yar_T-empty strain (Figure 1-C), indicating that its apparent interaction with the seven sRNA promoters could result from non-specific reporter activation. 29 TF-promoter interactions were thus retained, with 27 representing novel, previously unreported interactions. All results are summarised in Figure 1C, where the binding of TFs to sRNA promoters appears in red. In particular, SarA interacted with the promoters of *sprC* and *srn_9340*, consistent with previous studies (Figure S1-A) (21). On the other hand, several previously reported regulatory interactions were not recovered in our Y1H screen, in the case of CodY which is known to regulate *rsaD* expression, we have been unable to test its interaction on the *rsaD* promoter because we failed to introduce the pACTII-CodY plasmid into the Yar_011 (*PrsaD*) strain.

The 27 novel TF-promoter interactions identified in our Y1H screening provide a basis for exploring potential novel regulations. Notably, our screening results reveals that PyrR and ComK1 were capable of binding respectively the *rsaOI* and *rsaD* promoters. Moreover, ComK1 also binds the promoters of *sprC* and *srn_9340*. We can also observe that the promoters of *sprC* and *srn_9340*, whose regulation by SarA has been well described, share binding with 5 other TFs. This could be explained by their very similar promoter sequences, with 93% identity over the 29 bases located just upstream of the TSS (38). Finally, the promoter region of *srn_9340* was bound by 15 TFs. This may reflect a rather canonical promoter architecture that is permissive to TF binding in this heterologous system, although we cannot exclude that this sRNA is regulated by many TFs under highly variable conditions.

To visualize sRNAs sharing common regulatory characteristics, the results were integrated into a Venn diagram (Figure 1-D). Four TF-sRNAs groups emerge, such as NrdR with SprC and Srn_9340, SarA and HssR with SprC, Srn_9340 and sRNA71 or Comk1 with SprC, Srn_9340, and RsaD. VraR with SprC, Srn_9340 and Srn_9342. To investigate the regulatory interactions identified by Y1H screening in their native bacterial context, seven candidate TF–sRNA promoter interactions were selected for experimental validation in *S. aureus* using a β-lactamase reporter system (37). For each candidate, the same sRNA promoter as in the yeast assay was transcriptionally fused to the β-lactamase reporter, and promoter activity was compared between the wild-type strain and a strain lacking the TF of interest (Figure 1-E). Of the seven interactions tested, six were validated: ComK1 for *rsaD*, HssR for *sprC* and *sRNA71*, and SarA for *sRNA71* (Figure S1-B), together with VraR and MntR for *srn_9342*. For the latter two, β-lactamase activity increased by more than 2.5-fold in the absence of *vraR* or *mntR*, confirming their involvement in the regulation of *srn_9342* transcription and indicating that VraR and MntR act as repressors of *srn_9342* expression (Figure 1-F). Interestingly, VraR and MntR transcription factors were previously characterized for their regulatory roles on *rsaOI* and *rsaC*, respectively (33, 40). These observations suggest that Srn_9342 might participate in two distinct regulatory networks. The first network would link Srn_9342 with RsaOI through VraR regulation, while the second would connect Srn_9342 with RsaC via MntR. This dual regulatory potential is particularly intriguing given the known functions of these sRNAs: RsaOI mediates adaptation to cell wall stress induced by acidic conditions or vancomycin exposure, whereas RsaC contributes to ROS detoxification in manganese-rich environments by repressing *sodA* translation in favor of *sodM* expression (30).

Given these potential links between Srn_9342 and established stress response pathways, we have prioritized these three sRNAs, Srn_9342, RsaOI, and RsaC, for further investigation to elucidate whether and how they are functionally connected and to determine their coordinated regulatory roles in *S. aureus* physiology.

### Srn_9342 interacts with the stress-associated sRNAs RsaOI and RsaC

The potential interactions of Srn_9342 with both RsaOI and RsaC were first support by published MAPS data examining Srn_9342’s RNA partners, in which both RsaOI and RsaC independently enriched with Srn_9342 suggesting their interaction *in vivo* (35). Our findings prompted us to consider Srn_9342, RsaOI, and RsaC as a potential regulatory trio rather than studying them as separate pairs (Figure 2-A).

**Figure 2.**
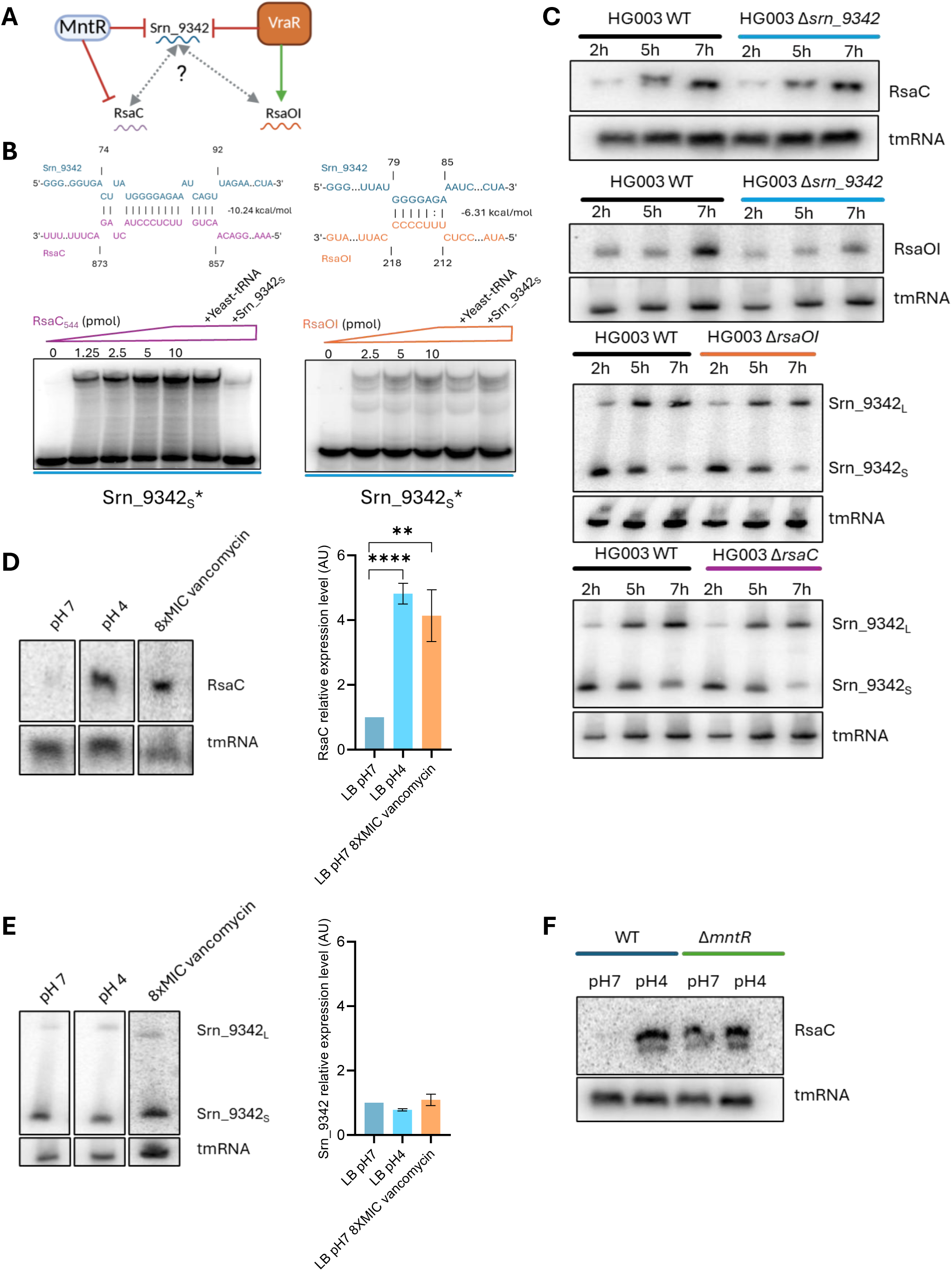
Regulatory network between Srn_9342, RsaOI, and RsaC: interaction validation and expression analysis. (**A**) Schematic of the proposed interaction network between Srn_9342, RsaC and RsaOI. (**B**) IntaRNA prediction of interaction and validation of Srn_9342-RsaOI and Srn_9342-RsaC interactions by EMSA. (**C**) Expression of *srn_9342*, *rsaOI*, and *rsaC* in HG003 WT, Δ*srn_9342*, Δ*rsaOI*, and Δ*rsaC* strains at different growth time points, analyzed by northern blot using *srn_9342*, *rsaOI*, or *rsaC*-specific probes, with tmRNA used as loading control. Cells were grown in TSB medium prior to the assay. (**D**) Expression of *rsaC* under acid and vancomycin stress conditions analyzed by northern blot and normalized to the control gene *tmRNA.* Cells were grown for 2 h in LB medium adjusted to pH 7, then pelleted and resuspended in either LB medium adjusted to pH 7, pH 4, or pH 7 supplemented with vancomycin at 16 µg/mL for 30 min prior RNA extraction. (**E**) Expression of *srn_9342* under acid and vancomycin stress conditions was analyzed by northern blot and normalized to the control gene *tmRNA.* Cells were grown and prepared as in (D). (**F**) Expression of *rsaC* in HG003 WT and Δ*mntR* strains was analyzed by Northern blot. Cells were grown in LB medium adjusted to pH7 for 2h and then exposed to either LB adjusted to pH7 or pH4 for 30 minutes prior to RNA extraction. Statistical analysis was conducted using Student’s *t*-test. Error bars represent the average of three independent experiments. Statistical significance is indicated by bars and asterisks as follows: \*\**P*< .01, \*\*\*\**P*<.0001

It’s important to note that Srn_9342 exists in two isoforms: a short 150 nucleotides (nt) form produced early in growth that is gradually replaced by a longer, 250 nt form during growth. At 5 hours of growth in rich medium, both isoforms of Srn_9342 are present in roughly equal levels. MAPS analysis showed that RsaOI was found enriched with the short form of Srn_9342, while RsaC was also enriched with both forms (35).

To further examine the association between RsaC and Srn_9342, we re-analyzed the published MAPS data using RsaC as a bait and created an annotated file allowing counting of Srn_9342 reads (Table S3) (30). This analysis revealed more than 4-fold enrichment of Srn_9342 providing additional support for an association between these two sRNAs. However, no RsaOI enrichment was detected in RsaC’s MAPS data.

To investigate direct interactions between Srn_9342 and RsaC or RsaOI, we first used IntaRNA to predict the sequences involved in duplex formation. The predicted interaction of Srn_9342 with RsaC involves 15 nucleotides between positions 74 and 92 of Srn_9342 (energy: -10.24 kcal/mol), whereas that with RsaOI involves nucleotides 79 to 85 (energy: -6.31 kcal/mol) (Figure 2B). As these two sites overlap, RsaC and RsaOI could bind Srn_9342 in a mutually exclusive manner or, alternatively, interact with it at different times or under different conditions. To confirm these *in silico* predictions, we assessed Srn_9342-RsaC and Srn_9342-RsaOI duplex formation using EMSA. We synthesized and radiolabeled an *in vitro* transcript corresponding to the short form of Srn_9342. Full length RsaOI was synthesized *in vitro*, while for RsaC, we focused on the predicted interaction region (nt 857-873) and synthesized both the full length of RsaC_1116_ and its truncated form lacking the repeated regions RsaC_544_ as previously described (30). As illustrated in Figure 2-B, the addition of RsaC resulted in the formation of a higher molecular weight complex whose intensity increased with rising RsaC concentrations. This interaction was specific, as evidenced by its competition with unlabeled Srn_9342 but not with non-specific tRNA. Similar results confirmed the specific interaction between Srn_9342 and RsaOI (Figure 2-B). We next examined interactions using the long form of Srn_9342. This isoform interacted with both RsaC_1116_ and RsaC_544_ (Figure S2-A), whereas an interaction with RsaOI was detectable but seems consistently weaker than with RsaC under *in vitro* condition tested (Figure S2-B). Together, these results show that Srn_9342 directly interacts with RsaC and RsaOI *in vitro*, potentially linking these sRNAs, although the functional relevance of these interactions *in vivo* remains to be determined.

### sRNA expression analysis across multiple stress conditions

To further investigate the regulation and potential functional relationships between Srn_9342, RsaOI, and RsaC, we analyzed their expression under different stress conditions and in relevant mutant strains. We extracted total RNA at three growth time points (2h, 5h, and 7h) from different strains (WT, *Δsrn_9342*, *ΔrsaC*, *ΔrsaOI*) and monitored three sRNAs (Srn_9342, RsaC, and RsaOI) by Northern blotting.

As shown in Figure 2-C, the absence of *srn_9342* did not detectably alter the expression profile of *rsaC* but appeared to slightly reduce the accumulation of RsaOI, this minimal and non-significative effect stayed unexplored. In parallel, we investigated the effect of *rsaC* and *rsaOI* deletion on Srn_9342 transcript level. Consistently with previous data obtained in TSB growth media, their deletion did not impact *srn_9342* expression pattern (41).

We next investigated whether the three sRNAs respond similarly to environmental stresses. Given that *rsaOI* expression under VraR control is strongly induced by acidic pH or vancomycin addition (33, 42), we investigated whether *rsaC* and *srn_9342* expressions were similarly induced under these stress conditions (Figure 2-D-E). The expression of *rsaC* was significantly induced at pH 4 and following treatment with 8×MIC vancomycin (Figure 2-D). In contrast, *srn_9342* expression levels remained unchanged under both stress conditions (Figure 2-E). To investigate whether VraR contributes to acid-induced expression of *rsaC*, we compared the RsaC levels in both WT or *vraR* deleted strains grown at pH 4 or pH 7. The results demonstrated that *rsaC* expression induction was similar regardless of *vraR* presence (Figure S3-A), indicating that VraR is not required for this induction. As MntR is a known repressor of *rsaC* expression (30), we further examined whether MntR influences its expression pattern. We observed pH-independent derepression of *rsaC* in the *mntR* mutant. The RsaC levels in the mutant strain at both pH were comparable to those in the wild-type strain at pH 4 (Figure 2-F).

Finally, manganese supplementation did not alter RsaOI or Srn_9342 levels (Figure S3-B-C), indicating that its effect is specific to *rsaC*. Srn_9342 levels were, in fact, stable across every condition tested in this study. Taken together, these results show that the three sRNAs studied have distinct expression patterns. While the abundance and temporal expression pattern of the two forms of Srn_9342 remained unchanged across the conditions tested, RsaC levels increased under acidic conditions and following vancomycin exposure. Unlike *rsaOI*, acid-induced *rsaC* expression did not require VraR. Furthermore, deletion of individual sRNAs had little effect on the abundance of the others.

### RsaOI and RsaC converge on the lac operon by targeting different genes

Since the expression of *rsaOI* and *rsaC* appears to respond to the same signals, however, by different regulatory mechanisms, we questioned whether the action of these two sRNAs in stress response is cooperative or redundant. To address this, we examined whether these two sRNAs share common mRNA targets.

Using the RNA predator software, which predicts sRNA-mRNA interactions in bacteria, we identified potential targets within a complete genome. These targets were ranked based on the energy required to disrupt the predicted interactions. To focus on the most robust targets, we set a threshold of interaction energy below -10 kJ/mol. A total of 432 and 535 potential targets met these criteria for RsaOI and RsaC, respectively, with 108 common targets. Functional gene annotations from the KEGG and NCBI databases were used to represent these candidates as a network (Figure S4). To identify potential functional redundancy or cooperation between RsaC and RsaOI, we focused on pathways in which both sRNAs could plausibly act together. Although *rsaC* is induced under manganese starvation (30), *rsaOI* expression is not affected by manganese availability (Figure S-3), making manganese homeostasis an unlikely context for cooperation between the two sRNAs. We therefore highlighted targets associated with galactose metabolism (outlined in green) and adaptation to vancomycin (outlined in orange). As expected, *lacD*, an established direct target of RsaOI (33), was found among the predicted RsaOI targets. Interestingly, the predicted RsaC targets did not include *lacD* but contained *lacC* mRNA (SAOUHSC_02453), which belongs to the same polycistronic mRNA. The predicted IntaRNA interaction is located at RBS of *lacC* involving nucleotides -12 to -4 upstream its start codon (Figure 3-A). Both genes belong to the *lacABCDFEG* operon, which encodes the tagatose-6-phosphate pathway for galactose utilization, *lacC* lying immediately upstream of *lacD* (Figure 3-A).

**Figure 3.**
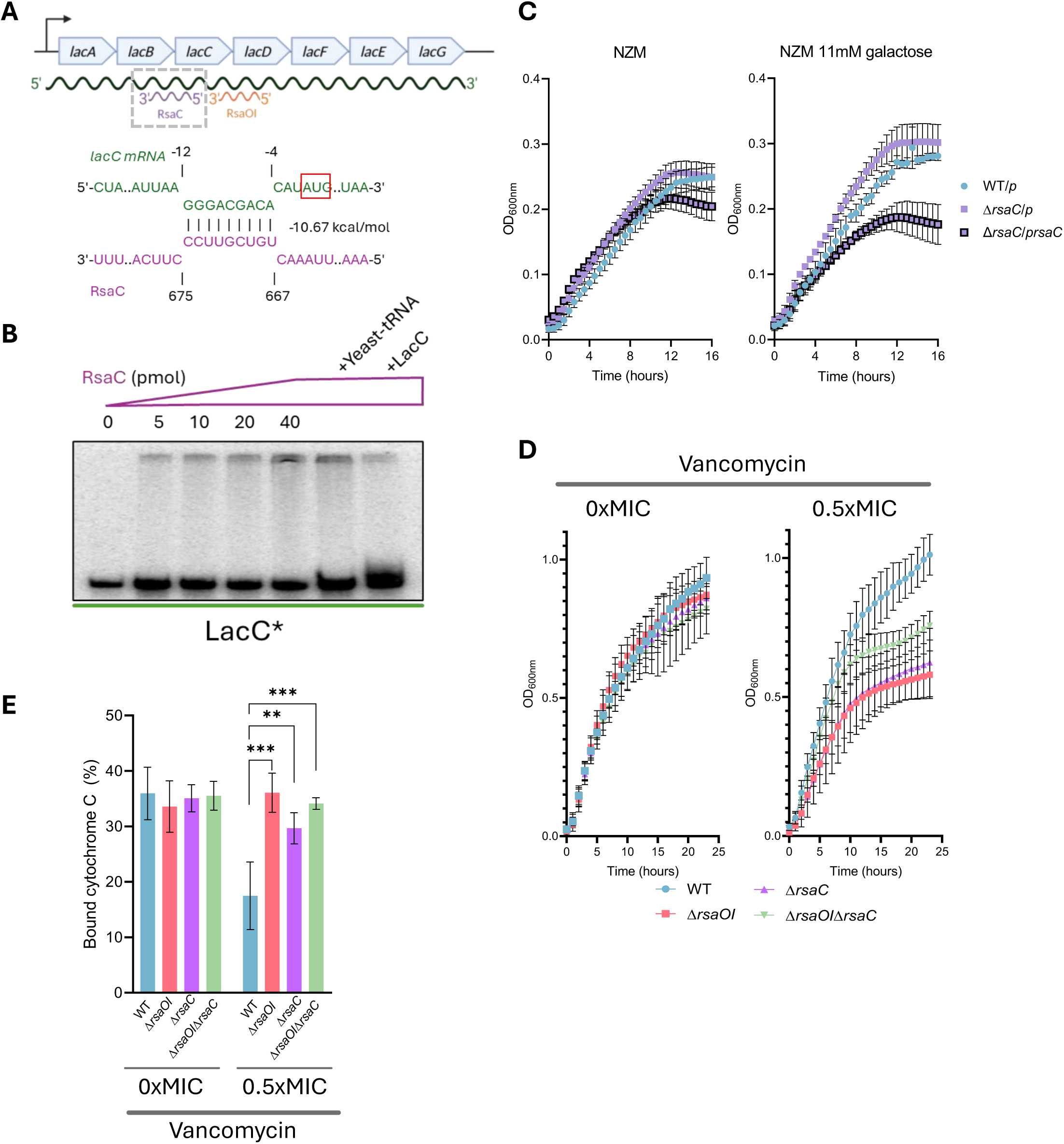
RsaC and RsaOI share *lac operon* as a common pathway and appear to contribute to the vancomycin response. (**A**) Schematic of the predicted RsaC–*lacC mRNA* interaction within the *lac* operon, together with the previously known RsaOI–*lacD mRNA* interaction (33) and IntaRNA prediction. (**B**) Validation of the RsaC–*lacC mRNA* interaction by EMSA. (**C**) Growth of HG003 WT, Δ*rsaC*, transformed with pICS3 (/*p*) or pICS3-*PamiA*-*rsaC* (/*prsaC*), in NZM medium supplemented or not with 11mM galactose. Growth was monitored using a Biotek microplate reader. (**D**) Growth of Δ*rsaOI*, Δ*rsaC*, and double-deletion strains in LB medium adjusted to pH7, with or without vancomycin at 1µg/mL (0.5xMIC). Growth was monitored using a Biotek microplate reader. (**E**) Cytochrome C binding assay in HG003 WT, Δ*rsaOI*, Δ*rsaC* and Δ*rsaOI*Δ*rsaC* strains. Cells were grown during 24h in LB medium adjusted to pH7 with or without exposure to vancomycin at 1µg/mL prior to the assay. Statistical analysis was conducted using Student’s *t*-test. Error bars represent the average of three independent experiments. Statistical significance is indicated by bars and asterisks as follows: \*\**P*< .01, \*\*\**P*< .001.

To experimentally test the *in silico*-predicted interaction between RsaC and *lacC* mRNA, we performed EMSA using the RsaC_544/1116_ constructs described above. The *lacC* mRNA which is normaly part of a polycistronic mRNA was synthesized *in vitro* from -113 nucleotides upstream from start codon to +116 nucleotides downstream start codon. When incubated with increasing concentrations of RsaC_544,_ we observed the formation of a higher molecular weight complex, indicating direct sRNA-mRNA interaction. To confirm the specificity of this interaction, we demonstrated that a non-specific competitor RNA did not disrupt the complex, while unlabeled *lacC* mRNA effectively competed with the radiolabeled mRNA, causing disruption of the complex (Figure 3-B). These findings validate the predicted binding of RsaC to *lacC*.

The involvement of RsaOI in galactose utilization has been previously reported (33). To determine whether RsaC also affects this pathway, we compared the growth of the wild-type, Δ*rsaC* and Δ*rsaC* overexpressing RsaC strains in NZM medium with or without galactose (Figure 3-C). In the presence of galactose supplementation, growth of both the Δ*rsaC* and wild-type strains were improved compared with the galactose-free condition, with the Δ*rsaC* strain growing slightly better than the wild type. In contrast, growth of the *rsaC*-overexpressing strain was not improved by galactose supplementation. These results parallel the phenotype described for RsaOI and suggest that RsaC impairs galactose utilization by inhibiting *lacC* translation, in line with the predicted interaction at its RBS. The two sRNAs thus appear to converge on the same operon by targeting different genes, rather than by sharing a single target.

### RsaOI and RsaC are required for surface charge remodeling under vancomycin stress

Because both RsaOI and RsaC are induced by vancomycin and by acidic pH, two stresses that affect the integrity of the cell wall (43, 44), we focused on the cell wall as a possible common target of their regulation. To assess the contribution of these sRNAs to vancomycin tolerance, we monitored growth kinetics in LB medium buffered at pH 7 at a subinhibitory concentration (0.5x MIC) that did not affect the growth of the WT strain. This assay quantifies the population growth rate over time rather than measuring population size at a single time point, and thus detects effects that endpoint colony-formation assays cannot resolve. As shown in Figure 3-D, WT, single and double Δ*rsaC* / Δ*rsaOI* mutants strains exhibited comparable growth in LB medium at pH 7, while vancomycin addition specifically impaired the growth of both single and double mutants without affecting the wild-type strain. Surprisingly, the Δ*rsaC*Δ*rsaOI* double mutant appeared slightly less affected by vancomycin than the single mutants, though this growth improvement did not reach statistical significance. The fact that the Δ*rsaC*Δ*rsaOI* double mutant growth is less affected by vancomycin (0.5xMIC) than single mutants suggests that RsaC and RsaOI may act on the cell wall stress response through both shared and independent pathways.

Given that vancomycin targets the bacterial cell wall, we investigated surface charge modifications using cytochrome c binding assays. This positively charged protein effectively reports on bacterial surface charge by binding to negative surface components (45), providing a sensitive indicator of cell wall adaptations. Our results (Figure 3-E) reveal a critical adaptive mechanism, WT strain significantly alter their surface charge in response to vancomycin stress (0.5×MIC), demonstrating an important survival strategy against antibiotic pressure (46). This adaptive response is completely abolished in all mutant strains (Δ*rsaC*, Δ*rsaOI*, and Δ*rsaC*Δ*rsaOI*), which maintain constant surface charges regardless of vancomycin exposure.

The inability of the mutant strains to modify their surface charge correlates with their impaired growth in vancomycin-containing medium (Figure 3-D), supporting a link between surface charge modification and the ability to grow in the presence of 0.5× MIC of vancomycin (46).

### Identification of RsaC and RsaOI targets in cell wall structure and remodeling

To identify potential targets of RsaC and RsaOI among genes involved in cell wall structure and remodeling, we selected genes encoding key enzymes acting at successive steps of cell wall biogenesis. These included *murD*, *murG* and *pbp2*, which catalyze, respectively, a cytoplasmic step and a membrane step of peptidoglycan precursor synthesis, and peptidoglycan cross-linking at the cell surface (47); *tagO* and *tarL*, which initiate and elongate wall teichoic acid (WTA) chains (47); *sscB*, involved in ssc polymer synthesis (48); *dltA*, required for D-alanylation of teichoic acids (49); and *mprF*, which lysinylates phosphatidylglycerol (50). The last two modify the net charge of the cell surface. We compared the expression of these genes in strains deleted for, or overexpressing, either sRNA. None of them (*murD*, *murG*, *pbp2*, *tagO*, *tarL*, *sscB*, *dltA* and *mprF*) showed expression differences across the strains tested. We also analyzed *spa*, encoding the cell wall associated protein A, and *SAOUHSC_02820*, encoding a putative lantibiotic transporter that may contribute to surface charge modulation under vancomycin stress (46). Both genes were oppositely affected by the two sRNAs (Figure 4-A), *rsaOI* overexpression decreased their expression, whereas *rsaC* overexpression increased it. Because sRNAs can act by blocking translation initiation without necessarily affecting transcript levels, the absence of changes in mRNA abundance for the other genes does not rule out the possibility of regulation at the protein level.

**Figure 4.**
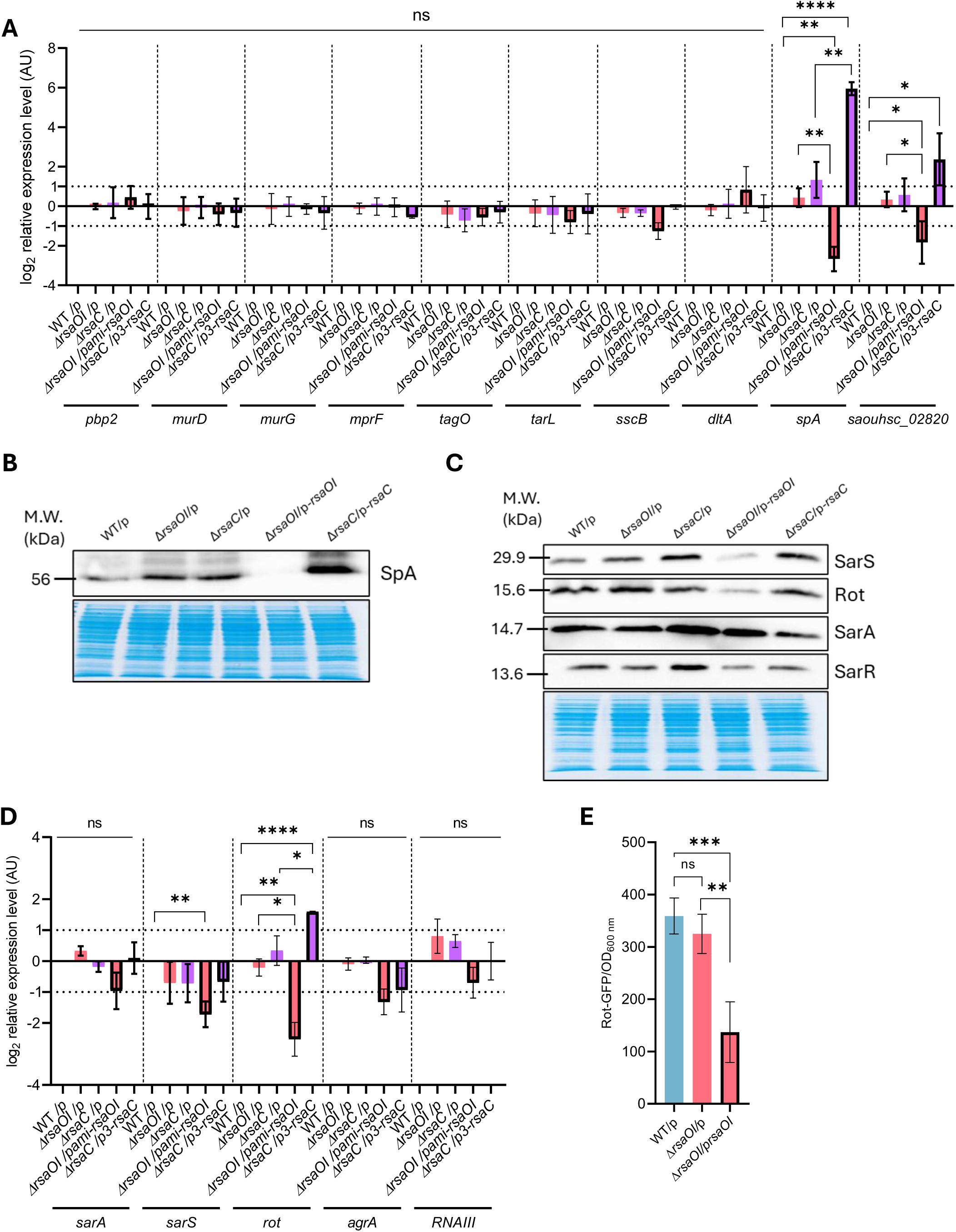
Effect of RsaOI and RsaC on cell wall gene expression and protein A production. (**A**) HG003 WT, Δ*rsaOI* and Δ*rsaC* were transformed with pICS3 empty vector *(/p*), pICS3*-Pami-rsaOI* (*/p-rsaOI*) or pCN51*-P3-rsaC* (/p3-rsaC), and expression of genes involved in cell wall metabolism was analyzed by RT-qPCR, normalized to the control gene *gyrB* using the 2^-ΔΔCt^ method. Cells were grown in LB medium adjusted to pH7 for 5h prior to RNA extraction. (**B**) Protein A expression levels were assessed by western blot on protein extracts from HG003 WT, Δ*rsaOI* and Δ*rsaC* strains transformed with pICS3-empty vector (/*p*), pICS3-*PamiA*-*rsaOI* (/*p-rsaOI*) and pICS3-*PamiA*-*rsaC* (/*p-rsaC*). Ready Blue staining of total protein extracts was done to compare loaded protein levels. Cells were grown as in (A) prior to protein extraction. (**C**) SarS, Rot, SarA and SarR expression level was analyzed by Western blot using the same strains as in (B). Cells were grown and prepared as in (A). (**D**) RT-qPCR analysis of *sarA*, *sarS*, *rot*, *agrA* and *RNAIII* expression under the same conditions as (A). (**E**) The effect of RsaOI on *rot* expression was analyzed using *gfp* translational reporter assay, using HG003 WT and Δ*rsaOI* strains co-transformed with pCN33-*PtufA*-*rot*-*gfp* and pICS3 empty vector *(/p*) or pICS3*-Pami-rsaOI* . The fluorescence intensity was measured after 3h of growth in LB medium adjusted to pH7. Statistical analysis was conducted using Student’s *t*-test. Error bars represent the average of three independent experiments. Statistical significance is indicated by bars and asterisks as follows: \**P*< .05, \*\**P*< .01, \*\*\**P*< .001, \*\*\*\**P*< .0001.

We then focused on protein A, a major virulence factor of *S. aureus*, and used Western blot to determine whether the changes in *spa* mRNA levels were reflected at the protein level (Figure 4-B). Protein A levels mirrored the mRNA data, it was undetectable upon *rsaOI* overexpression and increased upon *rsaC* overexpression. Because RsaOI and RsaC are naturally induced by acidic pH and vancomycin, we asked whether protein A levels also vary under these conditions. In the wild-type strain, protein A levels decreased markedly at pH 5 and under vancomycin (4× MIC) compared with pH 7 (Figure S5-A), when both sRNAs are upregulated.

Because both RsaC and RsaOI affected *spa* transcript and protein levels (Figure 4-A), possibly through an indirect mechanism, we monitored the amounts of SarS, Rot, SarA and SarR, which are known to control *spa* transcription, in strains deleted for or overexpressing *rsaOI* or *rsaC* (Figure 4-C). Deletion of *rsaOI* did not affect the level of any of the proteins tested, whereas its overexpression strongly decreased SarS and Rot levels. Overexpression of *rsaC* slightly decreased SarA, consistent with the increase in SarA observed in the *rsaC*-deleted strain. Such effect on SarA is consistent with previous evidence that *sarA* is a direct target of RsaC (30). Altogether, these results indicate that RsaOI reduces the amounts of SarS and Rot, two positive regulators of *spa*, in agreement with its negative effect on protein A production. In contrast, the effects of RsaC on these regulators were too weak to readily explain the increase in protein A levels, raising the possibility of a Rot- and SarS-independent mechanism.

We next analyzed whether *rsaOI* or *rsaC* overexpression altered the transcription of *sarA*, *sarS*, *rot*, *agrA* and *rnaIII*. *sarA, agrA* and *rnaIII* expression changes were statistically insignificant for either deletion of overexpression of *rsaOI* and r*saC*, (Figure 4-D). In the same time, *sarS* expression significantly decreased upon *rsaOI* overexpression, whereas *rot* expression showed the most dramatic changes, being strongly downregulated by *rsaOI* overexpression and upregulated by *rsaC* overexpression. This pattern precisely mirrored the opposing effects of the two sRNAs on Protein A abundance.

To further test this relationship, we examined protein A abundance in strains mutated for some of these regulators. However, protein A was undetectable in Δ*rot* and Δ*sarS* strains (Figure S5-B), confirming that both Rot and SarS are essential for *spa* expression but preventing us from determining whether RsaOI and RsaC regulate protein A through Rot or SarS. As shown on figure S5-C, the absence of *rnaIII*, *rnaIII and agrA* or *sarA* greatly increased SpA. Nevertheless, *rsaOI* overexpression dramatically reduced SpA production in these strains, whereas the positive effect of *rsaC* overexpression was difficult to observe probably because SpA is already highly expressed in these mutants (Figure S5-C). Interestingly, RNApredator analysis identified *rot* as a potential RsaOI target (Figures S4, S6-A), ranking immediately next to *lacD*, a previously validated RsaOI target (33). IntaRNA predictions further mapped the RsaOI–*rot* interaction to a region immediately downstream of the Shine-Dalgarno sequence (Figure S6-B), a location consistent with translational repression. To test this prediction, we used a rot-GFP translational fusion, which showed that *rsaOI* overexpression significantly reduced Rot expression at post transcriptional level (Figure 4-E), supporting a direct interaction. Moreover, *rsaOI* overexpression effect on rot-GFP translation is independent of *RNAIII* or *sarA* presence as shown on Figure S6-C.

Our results demonstrate that RsaOI and RsaC regulate protein A expression through distinct molecular mechanisms. Specifically, RsaOI and RsaC modulate *spa* expression at both the mRNA (Figure 4-A) and protein (Figure 4-B) levels, with opposite effects: *rsaOI* overexpression abolished protein A production, whereas *rsaC* overexpression increased it. This inverse regulation pattern suggests that these two sRNAs exert opposing effects on protein A production. The ability of RsaOI to pair with the *rot* mRNA *in silico* (Figure S6-C) is consistent with a direct post-transcriptional control, although this remains to be confirmed *in vivo*. In contrast, no such interaction was detected for RsaC, suggesting an indirect effect.

### RsaOI and RsaC impact *psma* expression

During protein gel electrophoresis and Coomassie staining, interestingly, we observed an intense low molecular weight band in *RNAIII* deletion strain that disappeared upon overexpression of either *rsaC* or *rsaOI* (Figure 5-A). Excised band was analyzed by mass spectrometry. As shown in Figure 5-A and Table S3, PSMα peptides 1-4 were identified in this band. These findings suggest that overexpression of RsaC and RsaOI may negatively impact PSMα production, as evidenced by the disappearance of the corresponding protein band in our experimental conditions.

**Figure 5.**
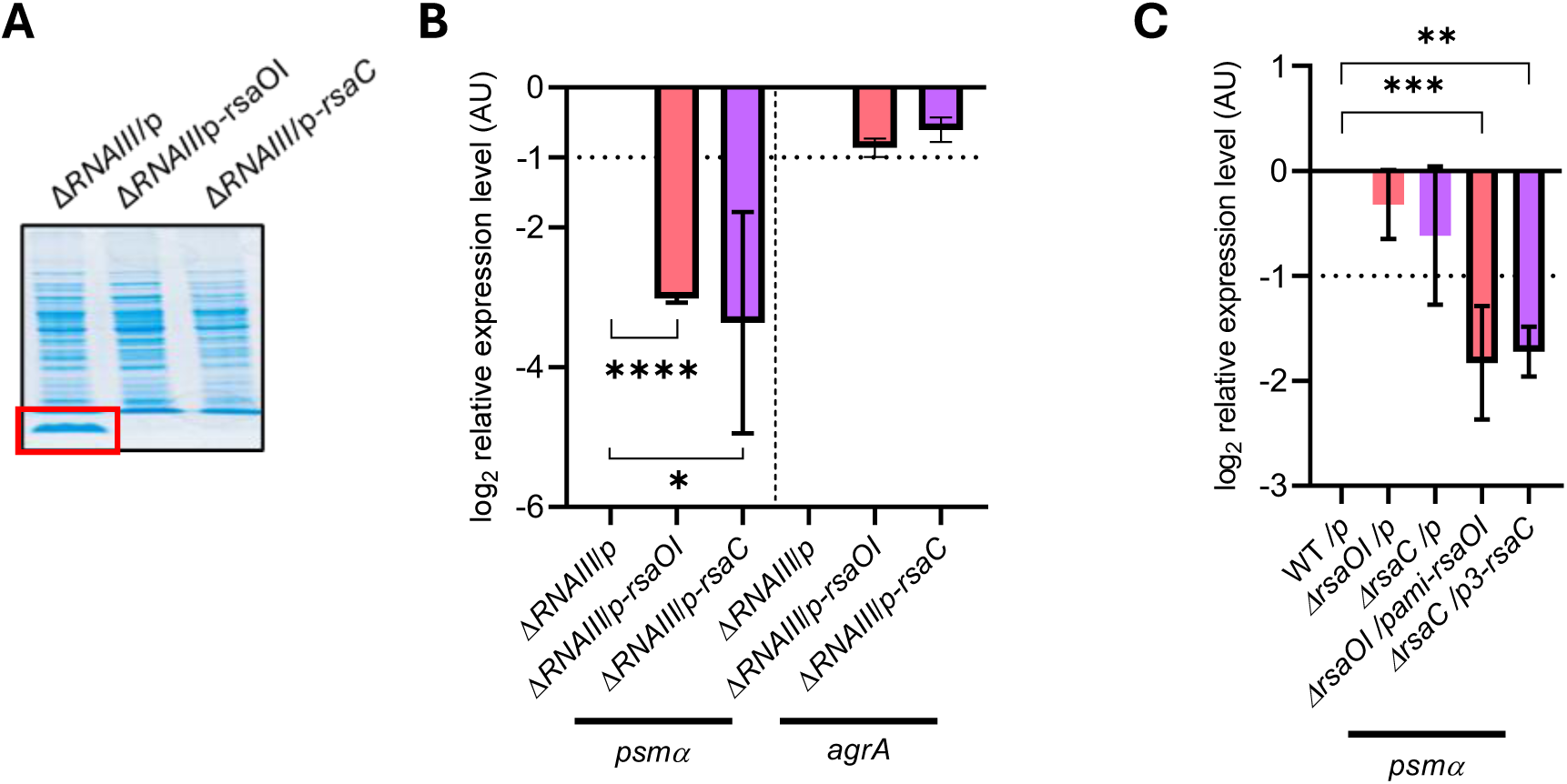
Effect of RsaOI and RsaC on PSMα production. (**A**) SDS-PAGE analysis of protein extracts from HG003 Δ*rnaIII* transformed with pICS3-empty vector (/*p*), pICS3-*PamiA*-*rsaOI* (/*prsaOI*) and pICS3-*PamiA*-*rsaC* (/*prsaC*). The band of interest is outlined in red. Cells were grown in LB medium adjusted to pH7 for 5h prior to protein extraction. (**B**) Using the same strains as in (A), the expression of genes involved in *psmα* synthesis and *agrA* were analyzed by qPCR, normalized to the control gene *gyrB* using the 2^-ΔΔCt^ method. Cells were grown as in (A) prior to RNA extraction. (**C**) HG003 WT, Δ*rsaOI* and Δ*rsaC* were transformed with pICS3 empty vector *(/p*), pICS3*-Pami-rsaOI* (*/p-rsaOI*) or pCN51*-P3-rsaC* (/p3-rsaC), cells were grown as in (A) prior to RNA extraction, expression of genes involved in *psmα* synthesis was analyzed by qPCR and normalized to the control gene *gyrB* using the 2^-ΔΔCt^ method. Statistical analysis was conducted using Student’s *t*-test. Error bars represent the average of three independent experiments. Statistical significance is indicated by bars and asterisks as follows: \**P*< .05, \*\**P*< .01, \*\*\**P*< .001, \*\*\*\**P*< .0001.

To test this hypothesis, we analyzed the expression of *psma* operon genes by RT-qPCR in a WT and a *rnaIII* deletion strain overexpressing or not *rsaOI* or *rsaC*. Consistent with the Western-blot results, we observed a decrease of *psma* expression related with *rsaOI* and *rsaC* overexpression (Figure 5-B-C).

Since AgrA is described as a positive regulator of *psma* expression, we studied its expression based on the overexpression of *rsaOI* or *rsaC*. According to Figure 5-B and Figure 4-D both in a WT and *rnaIII* deletion strains a non-significant decrease in *agrA* expression was observed after overexpressing *rsaOI* and *rsaC*. These results suggest that *rsaOI* and *rsaC* overexpression effect on *psma* does not involve *agrA* expression modification.

## Discussion

Small RNAs (sRNAs) are pivotal post-transcriptional regulators in *S. aureus*, coordinating adaptive responses to environmental stresses (24). Several sRNAs have been directly linked to both virulence and metabolism. Pro-virulence sRNAs, such as RNAIII, SprD, SprX and Teg41, promote the expression of virulence genes, whereas others, such as SprC, SprY and RsaA, repress it, either directly or indirectly (24). Here, we show that RsaOI and RsaC respond to the same cell wall stresses, including acidic pH and vancomycin exposure, and are further linked by their ability to interact with a third sRNA, Srn_9342. Both RsaOI and RsaC sense cell wall stress signals and trigger adaptive metabolic responses and immune evasion.

Although considerable efforts were made to identify the molecular targets of sRNA and their regulatory mechanisms, much less is known about how sRNA expression is regulated and how sRNA are integrated into broader gene regulatory networks. This work aimed to fill this gap by identifying transcriptional factors regulating sRNA expression and facilitating the classification of sRNAs and TFs into functional clusters. For the identification of transcriptional regulators, we performed a Y1H screening in *Saccharomyces cerevisiae* and assessed the direct binding of 43 regulatory proteins to the promoters of seven sRNAs. The Y1H screening allow the identification of direct binding *in vivo*. However, as the screening is done in a heterologous organism some interaction can be lost especially those involving regulators that must be modify post translationally. That was probably the case for VraR which failed to interact with the *rsaOI* promoter, despite the well-documented regulatory relationship between VraR and RsaOI (33). As a response regulator in a two-component system, VraR exists in both phosphorylated and non-phosphorylated states depending on activation signals. Similarly, MntR exhibited binding to the *srn_9342* promoter but not to *rsaC*. In *S. aureus*, it is established that in the presence of its Mn²⁺ ligand, MntR binds the *mntABC* operon promoter and represses its transcription, thereby inhibiting RsaC production (30). The absence of interaction between VraR and the *rsaOI* promoter, as well as between MntR and the *rsaC* promoter, reflects the dependence of these regulators on specific effectors, which may be absent or inadequate in the yeast. These loss of some interactions through false negatives was expected. Nevertheless, known interactions like SarA on promoters of *sprC* and *srn_9340* were recovered. Despite those limitations this Y1H screening revealed 27 previously uncharacterized potential interactions.

To confirm in the native host the protein-DNA interactions detected in yeast, we performed transcriptional reporter assays in *S. aureus* for seven candidate TF-sRNA pairs. Six were validated: transcription of the reporter gene driven by the *sprC*, *srn_9340* or *sRNA71* promoters was affected by the absence of SarA or HssR. Similarly, we confirmed the regulation of the *rsaD* promoter by ComK1, and of the *srn_9342* promoter by VraR and MntR (Figure 1-F; Figure S1-B). These results support the relevance of the Y1H screen for identifying functional regulators of sRNA promoters in *S. aureus*, although direct binding in the bacterium remains to be demonstrated.

Several independent lines of evidence drew our attention to Srn_9342–RsaOI–RsaC sRNAs. Both VraR and MntR, known to control *rsaOI* and *rsaC* respectively, were shown, in this study, to act as repressors of *srn_9342* expression. Furthermore, through EMSA studies, we confirmed direct interactions between Srn_9342 and both RsaC and RsaOI *in vitro* (Figure 2-B). Moreover, *in silico* predictions suggest that Srn_9342 may interact with RsaOI and RsaC through overlapping binding sites. This could imply interference between the two interactions, unless they occur at different times or under different conditions. Cross-deletion analysis showed that loss of one sRNA did not substantially affect the steady-state levels of the other two (Figure 2-C). These results suggest that these interactions likely do not confer major effects on sRNA stability. Srn_9342 exists as two isoforms, a short form expressed during early growth and a long form expressed at later stages, whose synthesis is sigma factor B-dependent (41). Because sigma factor B is a global stress response regulator activated by multiple environmental stresses (35), isoform production could provide a plausible link between environmental cues and the timing of Srn_9342 activity. We therefore hypothesize that the temporal ordering of Srn_9342–RsaOI and Srn_9342–RsaC interactions is governed by differential binding affinities of each sRNA for the two isoforms.

Given that these three sRNA are directly interconnected and their expression is regulated by common transcriptional regulators, we tested whether they respond to shared environmental stimuli. *rsaC* expression is known to be repressed by MntR in the presence of manganese. We therefore examined the expression of *rsaOI* and *srn_9342* under this condition, but neither sRNA was affected. *rsaOI* belongs to the cell wall stress (CWS) stimulon, and its VraR-dependent expression is strongly induced by acidic pH or vancomycin exposure (33, 42). Under these stress conditions, *rsaOI* and *rsaC* are co-induced, whereas *srn_9342* expression remains unchanged (Figure 2-D, Figure S3). *rsaC* induction occurs independently of VraR (Figure S3-A) whether this involve MntR or an alternative transcriptional activator remains to be determined. The invariance of *srn_9342* across all stresses tested is consistent with a sponge or chaperone sRNA, whose role is to modulate the availability of its partner sRNAs rather than to respond to the stress itself.

Acidic stress and vancomycin exposure represent major inducers of the CWS response, a coordinated physiological adaptation characterized by extensive remodeling of the cell wall (42, 43, 51–54). Consistent with a role in this process, Δ*rsaOI* and Δ*rsaC* mutants fail to modify their surface charge under CWS (Figure 3-D). RsaOI has been reported to repress *atl* and *hpr* directly, which encode components of cell wall autolysis and carbon transport, respectively, thereby linking vancomycin treatment to compromised cell wall turnover and reduced carbon metabolism (42). Because *rsaOI* and *rsaC* respond to the same signals through distinct regulatory mechanisms, we wondered whether their actions are redundant or cooperative, and found that they converge on carbon metabolism and more precisely on galactose utilization.

RsaOI is known to repress *lacD* (33), and we show here that RsaC binds *lacC* mRNA *in vitro*, at a site predicted to overlap its RBS (Figure 3-A). Galactose supplementation did not improve the growth of the *rsaC*-overexpressing strain, whereas it enhanced that of the wild-type strain (Figure 3-B). Thus, the two sRNAs converge on the tagatose pathway by targeting different genes of the same operon, rather than by sharing a single target, and their roles in galactose metabolism could be cooperative. This may help the bacterium counteract the cell wall stresses (Figure 6). One plausible route starts from the tagatose pathway being the primary means of galactose assimilation in *S. aureus*, generating glyceraldehyde-3-phosphate and dihydroxyacetone phosphate that fuel core metabolism and indirectly support cell wall maintenance. Repression of the pathway could redirect phosphoenolpyruvate (PEP), normally consumed during galactose uptake via the PTS LacFE system, towards MurA, which uses PEP to convert UDP-GlcNAc into UDP-MurNAc, a critical precursor for peptidoglycan biosynthesis (55, 56). In addition, we showed that both RsaOI and RsaC are required for surface charge remodeling. This effect could participate to the global cell wall stress response even if the mechanism needs to be further investigated.

**Figure 6.**
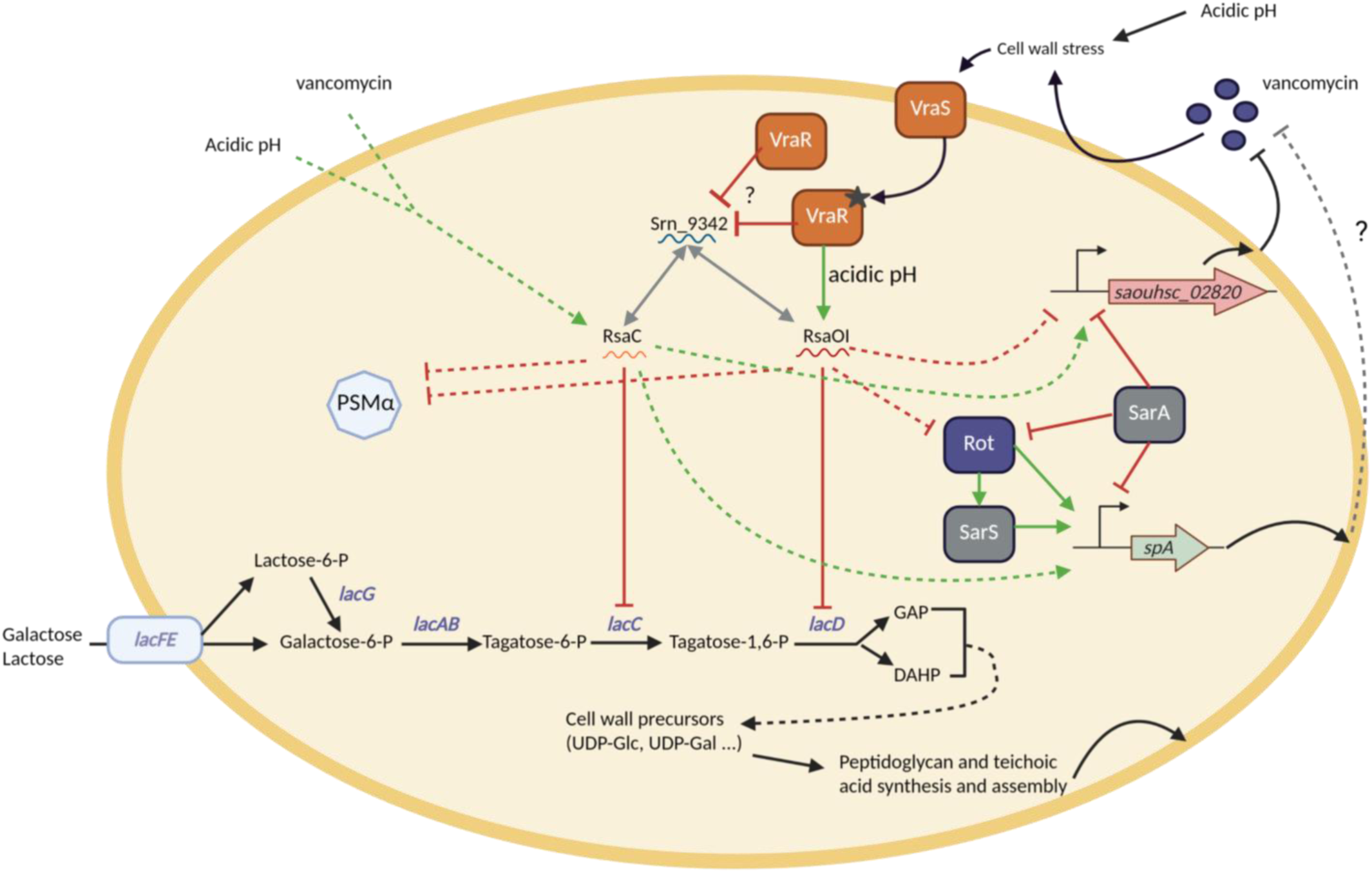
Proposed model of the RsaOI/RsaC regulatory network at the interface between cell wall stress response, galactose metabolism and virulence factor expression in *S. aureus*. Upon cell wall stress (vancomycin or acidic pH), the two-component system VraSR is activated and VraR directly activates *rsaOI* expression. *rsaC* expression is induced under the same conditions. Srn_9342 interacts with both RsaOI and RsaC. RsaOI and RsaC each repress one gene of the tagatose pathway (*lacD* and *lacC*, respectively), thereby limiting galactose utilization. Both sRNA also control *spa* expression indirectly and in opposite directions, possibly through Rot for RsaOI and SarA for RsaC. RsaOI and RsaC are further proposed to repress *psmα* production and to modulate *saouhsc_02820* expression. Solid lines indicate interactions supported by experimental evidence generated in this study. Dashed lines indicate predicted or not yet fully characterized interactions, arrows indicate activation and bars indicate repression.

Because sRNAs typically regulate multiple targets, we examined the expression of ten genes involved in peptidoglycan biosynthesis or cell wall structure. Two of them, *spa* and *saouhsc*_02820, responded oppositely to *rsaOI* and *rsaC* overexpression, *rsaOI* overexpression reduced their transcript levels, whereas *rsaC* overexpression increased them (Figure 4-A). For *spa*, protein levels mirrored transcript levels (Figure 4-B). As most sRNAs act post-transcriptionally by modulating translation or mRNA stability, a change in *spa* transcript level is unlikely to result from direct pairing and more plausibly involves an intermediate transcriptional regulator. The effects of RsaOI and RsaC on *spa* and *saouhsc_02820* are therefore likely indirect. Protein A, a cell wall-anchored protein, contributes to immune evasion by binding host IgG and limiting phagocytosis (57). Cell wall stress has been reported to reduce protein A production (58), and we observed lower protein A levels in the wild-type strain after acid or vancomycin exposure (Figure S5-A), conditions in which both sRNAs are induced. This is consistent with a repressive role for RsaOI under stress. How the opposite effect of RsaC fits into this response remains unclear and may depend on timing, as proposed in our model (Figure 6).

Protein A expression is controlled by several TFs and sRNAs, including RNAIII (59–61). At the protein level, *rsaOI* overexpression reduced Rot and SarS (Figure 4-C), two positive regulators of *spa* required for protein A production (Figure S5-B). RsaOI is predicted to pair with the *rot* mRNA at a site overlapping the ribosome-binding site (Figure S6-B), and it reduced the translation of a Rot-GFP reporter (Figure 4-E) independently of RNAIII and SarA (Figure S6-C). These data suggest that RsaOI limits protein A production by repressing Rot, possibly through direct pairing with *rot* mRNA, which remains to be demonstrated.

The situation is different for RsaC. No interaction with *rot* mRNA was predicted using the criteria applied to RsaOI, and the Rot protein level was not altered by *rsaC* overexpression, although *rot* transcript level increased (Figure 4-C-D). Independently of Rot, SarA is a possible intermediate between RsaC and protein A. RsaC pairs with the Shine-Dalgarno sequence of *sarA* mRNA and is expected to repress its translation (30), in agreement with the lower SarA level upon *rsaC* overexpression and its increase in Δ*rsaC* (Figure 4-C). Since SarA represses *spa* directly and through Rot (59, 62, 63), and protein A is strongly increased in a Δ*sarA* strain (Figure S5-C), reduced SarA could contribute to the effect of RsaC on protein A. These two routes are not mutually exclusive, and their respective contributions remain to be determined.

Overexpression of RsaOI or RsaC reduced the abundance of PSMα peptides, as well as *psmα* transcript levels (Figure 5; Figure S7). As for *spa*, we therefore hypothesize that these sRNAs act on PSMα indirectly, by targeting a transcriptional regulator of its expression. As *rsaOI* and *rsaC* overexpression impact PSMa production in the *RNAIII* deletion mutant (Figure 5-A-B), their effect on PSMa is RNAIII independent.

The PSMα family comprises short amphipathic peptides of 20–25 amino acids that exhibit potent cytolytic activity and modulate host inflammatory responses through neutrophil chemotaxis (64). Their balanced expression is important for tuning the trade-off between immune evasion and immune activation, particularly under stress conditions, which makes their repression during CWS consistent with the transient downregulation of Protein A described above.

Placing our results alongside published work, the most striking feature of this network is how many sRNAs converge on the same two virulence factors. At least five, RNAIII, RsaOI, RsaC, Teg41 and RsaE, modulate PSMα expression, and four of them (RNAIII, RsaOI, RsaC and Teg41) also influence Protein A regulation. Rather than adding two more sRNA–target pairs to a catalogue, our data therefore identify Protein A and PSMα as a tightly coordinated regulatory node under CWS, approached by several sRNAs through distinct mechanisms. Regarding protein regulators, *psmα* expression is governed by MgrA, (16, 65), SarA (66) and AgrA (15). Among the sRNAs, RNAIII represses *psmα* expression (67), while Teg41 and RsaE act as activators (68, 69). Teg41’s transcriptional effect appears to be indirect (65, 70) whereas RsaE, directly inhibit translation (69).

Two further observations reinforce this view of a coordinated node. Teg41’s involvement extends beyond *psmα* to *spa* expression, and RNA-seq-based network analysis positioned Teg41 within a stress-responsive module of 34 sRNAs that includes RsaOI, consistent with a role in antibiotic stress adaptation (71). The regulation of RsaE by the sponge RsaI provides a precedent for the architecture we propose for Srn_9342, which could titrate RsaOI and RsaC.

Our findings, are summarized in Figure 6. These data support a hypothesis in which Srn_9342 could titrate RsaOI and RsaC in a temporally controlled manner. Each sRNA may bind Srn_9342 alternatively, so that they may be recruited sequentially rather than together, progressively remodeling metabolic pathways and thereby modulating bacterial surface charge. We propose in this model that exposure to CWS, acidic pH or vancomycin leads to transcriptional induction of *rsaOI* and *rsaC*. RsaOI is supposed to act first, indirectly repressing both PSMα peptides and Protein A production by inhibiting *rot* mRNA translation and so limiting Rot-dependent activation of *spa* transcription. RsaC subsequently assumes regulatory control, maintaining repression of the tagatose pathway and of *psmα* expression. Because RsaC has a positive effect on *spa* expression, high cell density, reflecting successful proliferation, may relieve Srn_9342-mediated sequestration of RsaC and permit *spa* re-expression for immune evasion, while PSMa production is restored at this advanced growth phase through alternative regulators such as Teg41 or RsaE.

Together, our data identify RsaOI and RsaC as a regulatory module acting on cell wall stress, in which both sRNA converge on galactose metabolism and on the virulence factors Protein A and PSMα through distinct routes.

## Materials and methods

### Strains, plasmids and growth conditions

All strains and plasmids used in this study are listed in Table S1. *Escherichia coli* strains were grown in lysogeny broth (LB, MP Biomedicals) at 37 °C with shaking (160 rpm), or on LB agar at 37°C. *Staphylococcus aureus* strains were grown at 37 °C with shaking (160 rpm) in LB, brain heart infusion (BHI), tryptic soy broth (TSB) or NZM (Fluka) broth, depending on the assay. When required, LB was buffered to pH 7 with 100 mM HEPES and HCl. For plasmid or resistance-cassette maintenance, media were supplemented with antibiotics at the following concentrations: ampicillin, 50 µg/ml and kanamycin, 50 µg/ml for *E. coli*; chloramphenicol, erythromycin and tetracycline, 10 µg/ml and kanamycin, 250 µg/ml for *S. aureus*. Purified plasmids were introduced into *S. aureus* RN4220 by electroporation, purified and were then used to transformed HG003 or its derivatives.

To induce acid stress, overnights cultures were diluted in LB pH 7 to OD_600nm_ 0.1, then cells were grown for 2h in LB supplemented with 100 mM HEPES to maintain the pH at 7. Cultures were then split, pelleted by centrifugation (4500 rpm, 10 min) and resuspended for 30 min in fresh LB buffered with HEPES to pH 4 or pH 7. For metabolic assays, overnight cultures were diluted 1:100 in fresh LB or NZM broth and grown at 37 °C; where indicated, media were supplemented or not with galactose (11 mM). Growth was monitored with a Biotek microplate reader.

*Saccharomyces cerevisiae* strains were grown at 28 °C with shaking (160 rpm) in yeast extract–peptone–dextrose medium (YPD, MP Biomedicals) supplemented with adenine (6x10⁻⁵ M), or in yeast nitrogen base medium (YNB, MP Biomedicals) supplemented with glucose (20 g/l), adenine (6x10⁻⁵ M), tryptophan (10⁻⁴ M) and lysine (2.7x10⁻² M). When necessary, leucine (4.5x10⁻⁴ M) and histidine (1.3x10⁻³ M) were added.

### Yeast one hybrid screening

Yeast reporter strains were constructed by integrating the pYI2267OZ plasmid (72), previously linearized at its *Xho*I site, at the *URA3* locus of *S. cerevisiae* strain YM954. This plasmid carries the promoter of the sRNA of interest in transcriptional fusion with the *HIS3* gene, and its integration into the yeast genome confers uracil prototrophy. In parallel, a library of 43 pACTII-TF plasmids was generated by cloning the coding sequences of the TFs of interest (from start to stop codon) into the *Bam*HI and *Nco*I sites. Each plasmid carries a translational fusion between the *S. cerevisiae* GAL4_AD_ domain and the gene encoding one *S. aureus* TF of interest (listed in Table S1). Reporter strains were then transformed individually with the pACTII-TF plasmids, and transformants were selected on YNB without leucine (YNB-Leu).

Overnight cultures grown in YNB-Leu were adjusted to an OD600 of 1 and serially diluted 1:10 in sterile water. Aliquots of 8 µL of the 10⁰ to 10⁻³ dilutions were spotted onto YNB without leucine and onto YNB without leucine and histidine. Plates were incubated at 28 °C for 48 h before the results were recorded and photographed.

### Reporter Gene experiments

β-lactamase and GFP reporter assays were performed as previously described before (38, 73) respectively, with the following specifications. For β-lactamase assays, overnight cultures of *S. aureus* strains carrying pCN41c derivatives, with or without the sRNA promoters of interest (Table S1), were used to inoculate fresh cultures at an OD600_nm_ of 0.1. At each sampling time, bacteria were harvested by centrifugation and resuspended in a volume of PBS adjusted to normalize cell numbers to the OD600_nm_ measured at the time of sampling. Cells were lysed for 20 min at 37 °C in the presence of lysostaphin (0.7 mg/mL), benzonase (0.2 U/µL) and MgCl2 (0.1 mM). Nitrocefin was then added (0.25 mg/mL), and β-lactamase activity was quantified by measuring the OD at 492 nm every 10 min for 40 min at 37 °C in a Biotek microplate reader (Agilent, Synergy2 software). β-lactamase activity was normalized to the protein content of the same samples, determined in parallel by Bradford assay.

For GFP assays, a two-plasmid system was used: *S. aureus* strains carried pICS3-empty, pICS3-P*amiA*-*rsaOI* or pICS3-P*amiA*-*rsaC* together with pCN38-P*tufA*-*gfp*, pCN38-P*tufA*-*rot*-*gfp* or pCN38-P*tufA*-*lacC*-*gfp* (Table S1). Overnight cultures were diluted 1:100 in LB medium (pH 7), and fluorescence and OD600 were monitored in a Biotek microplate reader.

### RNA extraction, northern blots and RT-qPCR

Cells were collected and centrifuged for 10 min at 4500 rpm and 4 °C, then pellet were resuspended in RNA lysis buffer containing 0.5% SDS, 20 mM sodium acetate and 1 mM EDTA (pH 5.5). Total RNA extraction and northern blot assays were performed as previously described (38, 74). The DNA probes used for RNA detection are listed in Table S2. For sRNA detection, 10-20 µg of total RNA were separated on 8% polyacrylamide–8 M urea gels or 1% agarose formaldehyde gel for RNA>500nt and transferred onto Hybond-N+ membranes (Amersham). For mRNA detection, 10-20 µg of total RNA were separated on 1% agarose–formaldehyde gels and transferred onto Hybond-N+ membranes (Amersham). Membranes were hybridized with specific ³²P-labelled probes (Table S2) in ExpressHyb solution (Ozyme) and washed according to the manufacturer’s recommendations. They were then exposed and scanned with a Storm 825 (GE Healthcare), and image quantification was performed with ImageQuant Tool 7.0. Loading controls consisted of radiolabeled transfer-messenger RNA (tmRNA).

For reverse transcription quantitative PCR (RT-qPCR), total RNA samples were treated with the DNase I Amplification Grade Kit (Invitrogen) to remove residual DNA. cDNA synthesis and RT-qPCR were performed as previously described (73), using *gyrB* as the normalization control.

### *In vitro* transcription, RNA labeling and gel retardation assay

DNA templates were amplified from *S. aureus* HG003 genomic DNA by PCR using specific primers (Table S2), with forward primers carrying a T7 promoter sequence. RNAs were transcribed from these PCR products with the MEGAscript T7 kit (Ambion) and labelled at their 5′ end using [γ-³²P] ATP (Amersham Biosciences) and T4 polynucleotide kinase (Invitrogen). Binding reaction containing labeled and unlabeled RNAs were carried out in a final volume of 10 µL, and gel-shift assays were performed as previously described (33).

### Cell surface charge

Cell surface charge was estimated by cytochrome c binding, as previously described (45). Overnight cultures of *S. aureus* grown in LB buffered to pH 7, with or without vancomycin (1 µg/ml), were centrifuged, washed twice in 20 mM MOPS and resuspended in 20 mM MOPS to equalize the OD600_nm_ of all samples. Cytochrome c (0.5 mg/ml in 20 mM MOPS) was added to 200 µL of cell suspension. After 10 min of incubation at room temperature, the mixtures were centrifuged and the supernatants transferred to a 96-well microplate. The OD at 405 nm was measured with a Biotek microplate reader (Agilent).

### Total protein purification, western blots

Total proteins were extracted from *S. aureus* pellets, generally obtained from 25 ml cultures after 5 h of growth, using B-PER reagent (Thermo Fisher) according to the manufacturer’s instructions, supplemented with protease inhibitors (1X, Merck) and lysostaphin (10 µg/ml, Sigma), with an additional mechanical lysis step with glass beads in a FastPrep system. Lysates were centrifuged, and the supernatants containing total proteins were quantified with a Qubit fluorometer (Thermo Fisher) and stored at −20 °C. Equal amounts of protein were separated by SDS-PAGE at 120 V in Tris-glycine-SDS buffer, for a time adapted to the size of the protein of interest, and transferred onto 0.2 µm nitrocellulose membranes (Amersham) in Tris-glycine-ethanol buffer for 1 h 15 min at 40 mA. Membranes were washed and blocked in TBS–0.1% Tween supplemented with 5% skimmed milk, then incubated with the primary antibody for 1 h 15 min at 4 °C, followed by a horseradish peroxidase (HRP)-conjugated secondary antibody, with washes in TBS–0.1% Tween between each step and before detection. Proteins were detected with the ECL Prime Western Blotting detection kit (Amersham) and imaged with an ImageQuant LAS 4000 system. As a loading control, duplicate gels were stained with Ready Blue (Sigma) according to the manufacturer’s instructions.

### Identification of peptides by mass spectrometry

Total protein extracts (25–50 µg, prepared as described above) were separated on a 10% (37.5:1) polyacrylamide gel in Tris-glycine-SDS buffer, stained with Ready Blue (Sigma) and destained in water. The band migrating at ∼2.5 kDa was excised with a clean scalpel under keratin-free conditions, cut into pieces and stored at 4 °C until shipment. Protein identification was performed by the Proteom’IC facility (Institut Cochin, Paris, France). Gel pieces were digested with trypsin, and the resulting peptides were analyzed by nanoLC–MS/MS on a timsTOF HT instrument (Bruker) operated in data-dependent acquisition mode with a 30 min gradient. Spectra were searched with Mascot against the SwissProt database restricted to bacteria, with trypsin specificity, up to one missed cleavage, carbamidomethylation of cysteine and oxidation of methionine as variable modifications, monoisotopic masses, a precursor mass tolerance of 20 ppm and a fragment mass tolerance of 20 mmu. Identifications were retained above a significance threshold of *P* < 0.05 and an ion score cut-off of 25.

## ACKNOWLEDGEMENTS

We would like to thank our lab colleagues and more particularly N. Germain-Amiot for technical help. We are also grateful to P. Bouloc and W. Liu for strains. M. Hallier, for the Sar antibodies, strains and plasmids. L. Kuras for yeast strain and plasmid. C. Broussard and the Proteom’IC facility for PSM identification. I. Caldelari and P. Romby for plasmids, advices and for fruitful discussions.

This study was supported by Institut National de la Santé et de la Recherche Médicale INSERM, University of Rennes and UFR de Pharmacie de Rennes (to H.R.).

Investigation, methodology, visualization: H.R.; M.G.; C.S.; S.C.; A.R.; Resource: S.D.; Conceptualization and supervision: Y.A.; S.C. and A.R.; Manuscript writing: all authors.

We declare that we have no conflict of interest.

---

**Figure S1.**
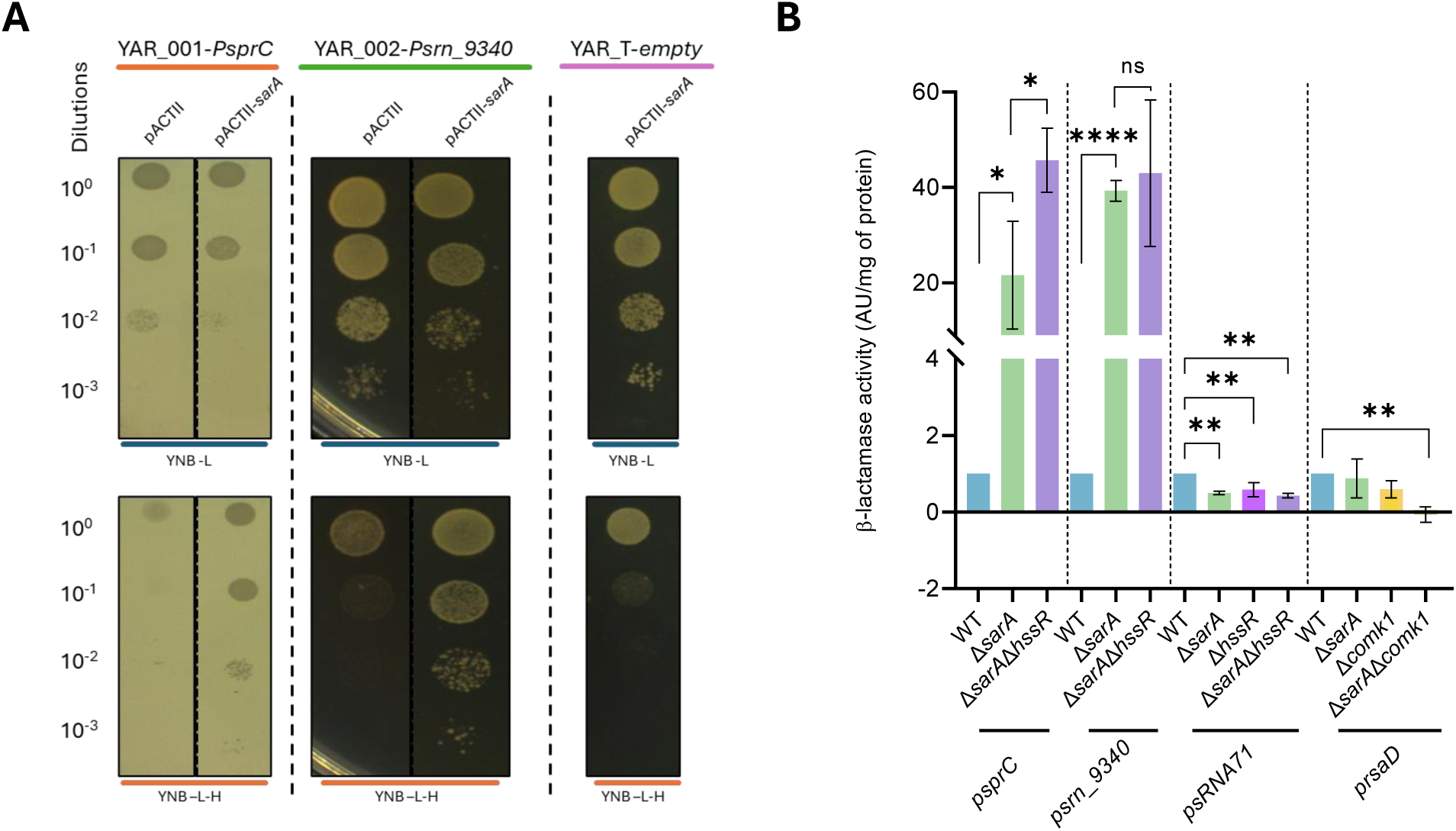
Validation of the Y1H TF-promoter interactions. (**A**) Recovery of previously known TF-promoter interactions. Serial dilution spot assay (10⁻³ to 10⁰) of the YAR_001-*PsprC*, YAR_002-*Psrn_9340*, and YAR_T-*empty* strains, transformed with pACTII or pACTII-*sarA*, on YNB medium supplemented or not with histidine, used to assess histidine prototrophy as a readout of Gal4AD-SarA binding to the tested promoters. (**B**) Validation in *S. aureus* of candidates TFs, on the *sprC, srn_9340*, *sRNA71* and *rsaD* promoters using the β-lactamase reporter assay. Cells were grown for 5h in TSB medium prior to the assay. Statistical analysis was conducted using Student’s *t*-test. Error bars represent the average of three independent experiments. Statistical significance is indicated by bars and asterisks as follows: \**P*< .05, \*\**P*< .01,\*\*\*\**P*< .0001.

**Figure S2.**
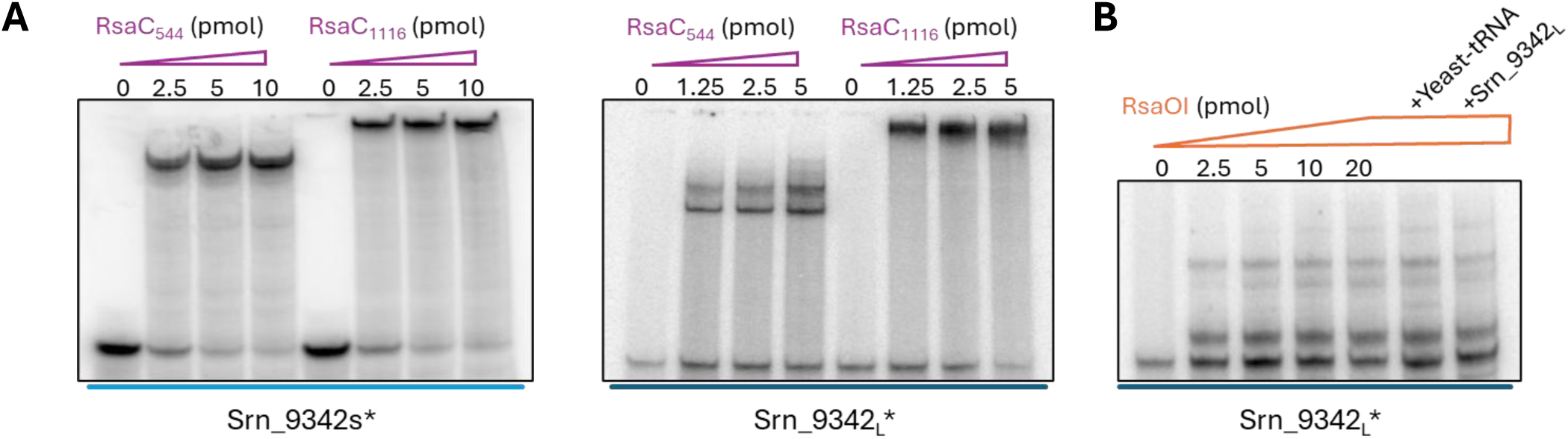
Srn_9342 interact with bot RsaC and RsaOI. (**A**) Validation of Srn_9342 with truncated RsaC_544_ and native RsaC_1116_ interactions by EMSA. (**B**) Validation of Srn_9342_L_ and RsaOI interaction by EMSA.

**Figure S3.**
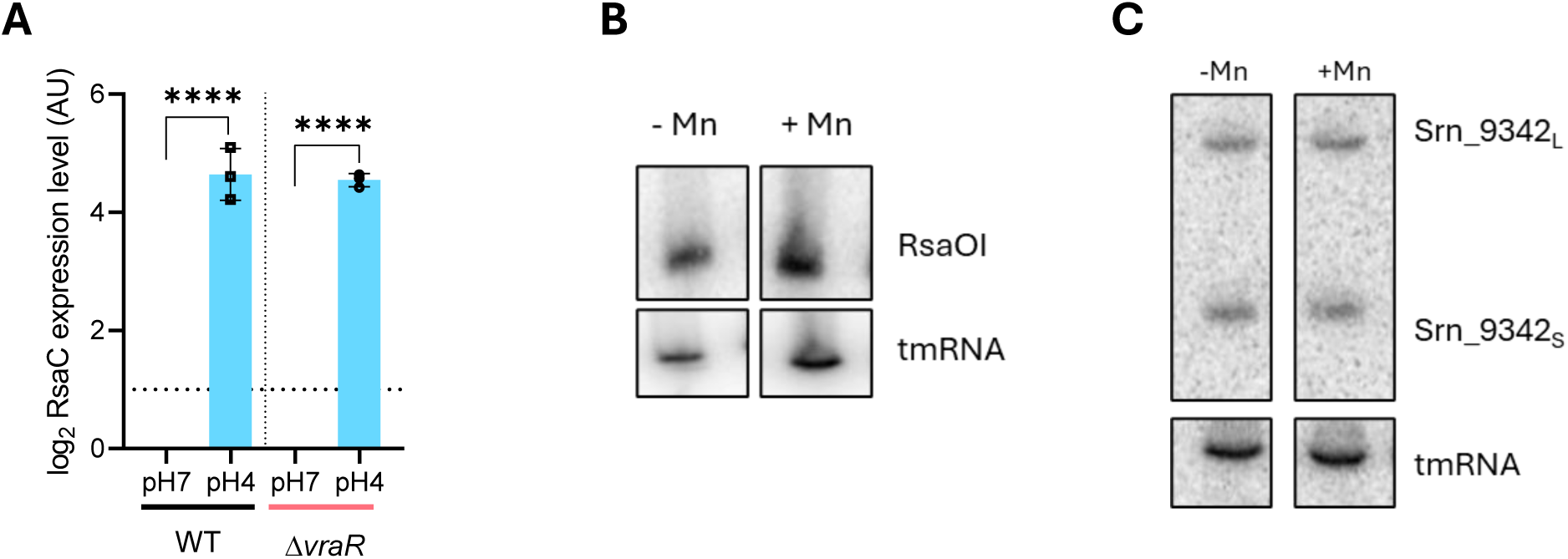
Expression of *rsaC*, *rsaOI* and *srn_3342* in response to pH or manganese availability. (**A**) Expression of *rsaC* in HG003 WT and Δ*vraR* strains was analyzed by RT-qPCR, normalized to the control gene *gyrB* using the 2^-ΔΔCt^ method. Cells were grown in LB medium adjusted to pH7 for 2h and then exposed to either LB adjusted to pH7 or pH4 for 30 minutes prior to RNA extraction. (**B**) Expression of *rsaOI* in HG003 WT was analyzed by Northern blot with tmRNA used as loading control. Cells were grown in LB medium for 5h supplemented or not with 25 µM MnCl_2_ prior RNA extraction. (**C**) Expression of *srn_9342* in HG003 WT strain was analyzed by Northern blot with tmRNA used as loading control. Cells were grown as in (B). Statistical analysis was conducted using Student’s *t*-test. Error bars represent the average of three independent experiments. Statistical significance is indicated by bars and asterisks as follows: \*\*\*\**P*< .0001.

**Figure S4.**
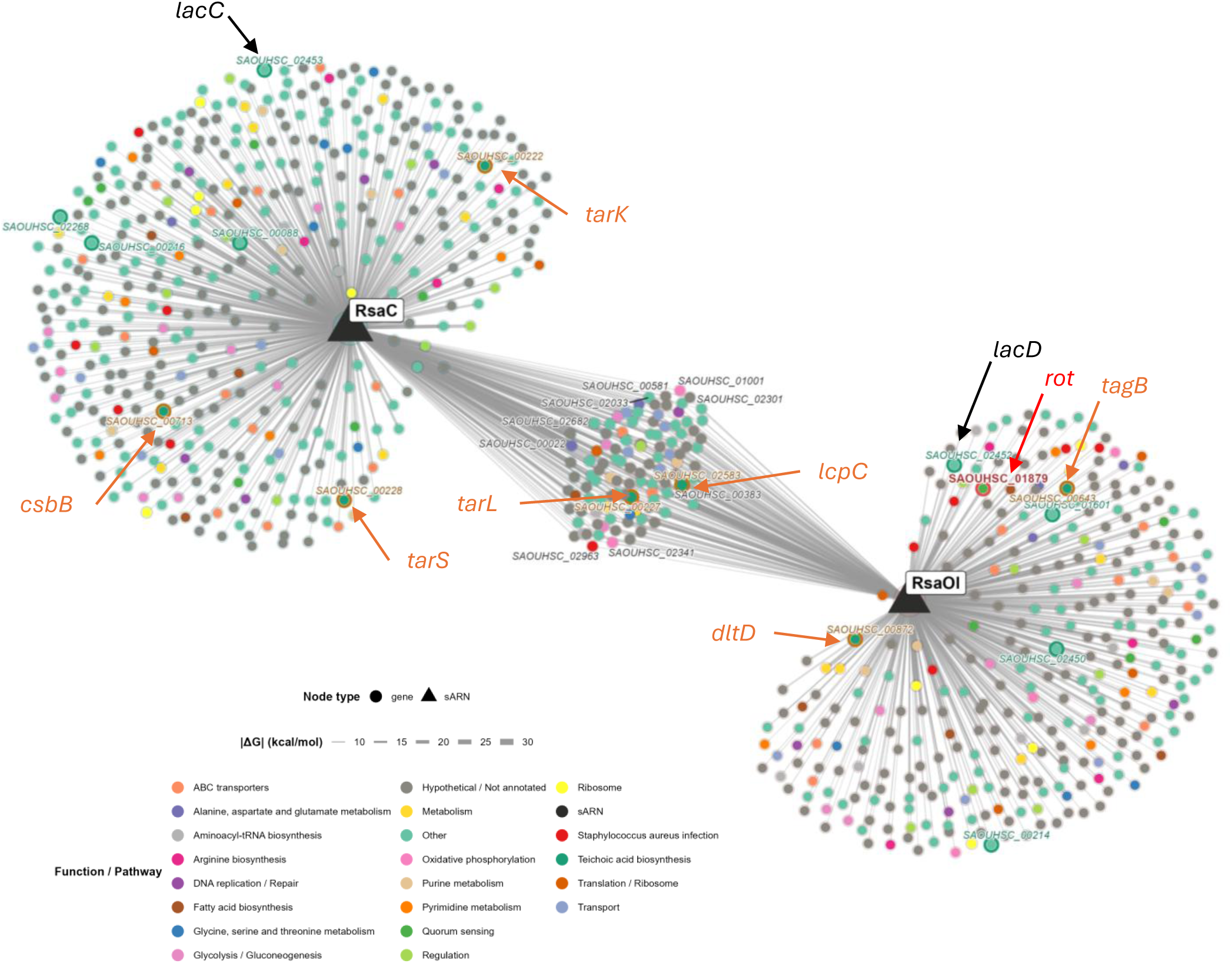
Predicted target network of RsaC and RsaOI reveals shared targets involved in cell wall stress response and galactose metabolism. Interaction network of RsaC and RsaOI predicted targets in *S. aureus* NCTC 8325, generated using RNApredator with an energy threshold of ΔG ≤ -10 kcal/mol. Triangular nodes represent the sRNAs (RsaC, RsaOI). Circular nodes represent predicted target genes, positioned according to a force-directed layout and colored according to their functional category (KEGG pathway annotation, or GFF-based functional category when no KEGG pathway was available). Node size is proportional to node degree, and edge width/transparency reflect the predicted interaction energy (|ΔG|). Genes predicted as targets of both RsaC and RsaOI are positioned in the central overlapping region and labeled with their locus tag. Genes involved in galactose metabolism, including *lacC*, are highlighted with a green halo. A TF, the repressor of toxin, *rot* (SAOUHSC_01879) is highlighted with a red halo. And genes involved in teichoic acid biosynthesis are highlighted with an orange halo.

**Figure S5.**
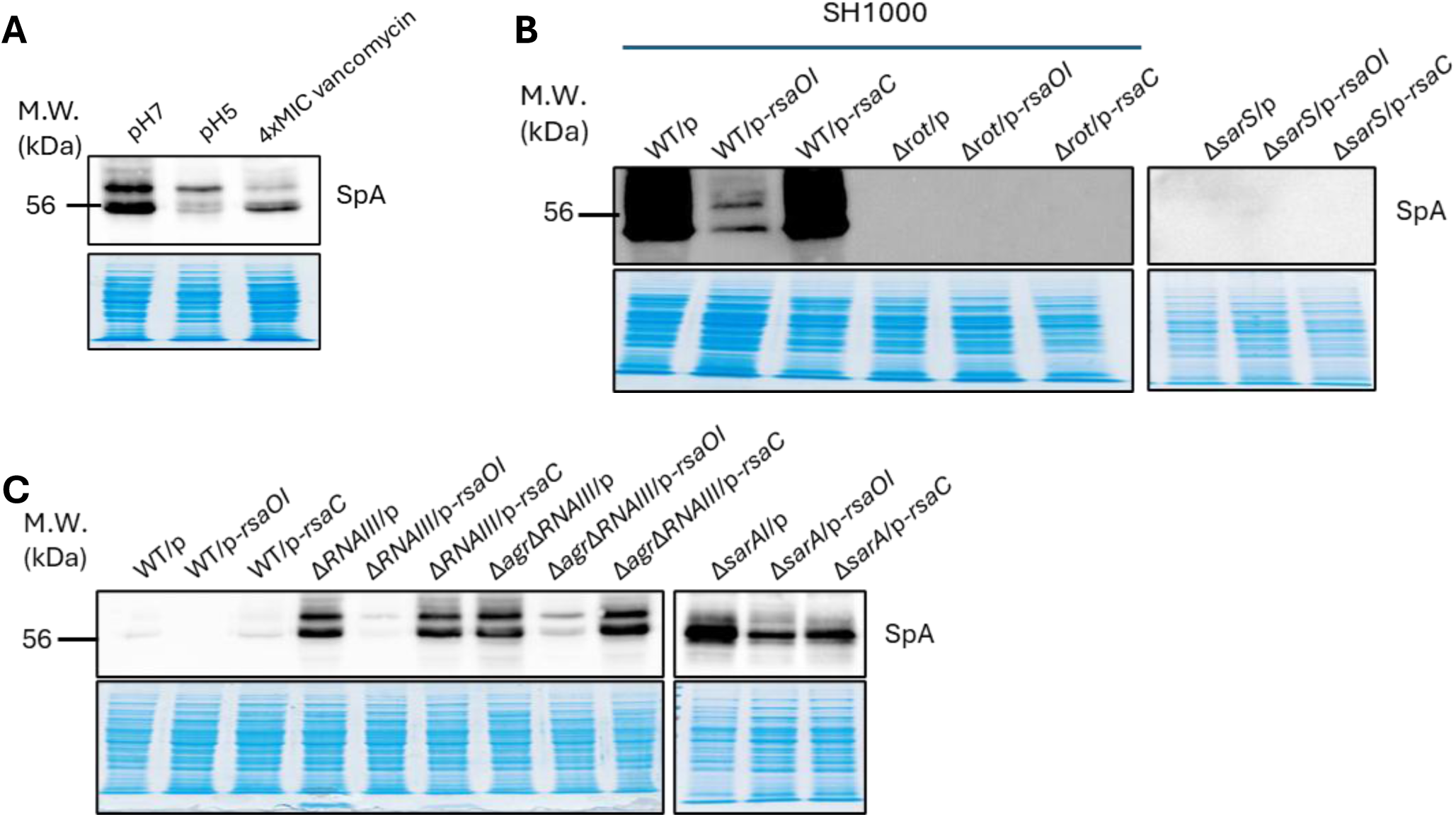
Protein A expression among different condition or in different strains overexpressing *rsaOI* or *rsaC*. (**A**) Protein A expression was analyzed by Western blot in HG003 WT strain grown for 3 h in LB medium adjusted to pH7 and then induced for 2 h in LB medium adjusted to pH 7, pH 5, or pH7 supplemented with vancomycin at 4xMIC (8 µg/mL). Ready Blue staining of total protein extracts was used as loading control. (**B**) Protein A expression was analyzed by Western blot in SH1000 WT, Δ*rot* and HG003 Δ*sarS* strains, transformed with the pICS3 empty vector (/p), pICS3-*Pami*-*rsaOI* (/p-*rsaOI*), or pICS3-*Pami*-*rsaC* (/p-*rsaC*). Ready Blue staining of total protein extracts was used as loading control. (**C**) Protein A expression was analyzed by Western blot in HG003 WT, Δ*RNAIII*, Δ*RNAIII*Δ*agr* and Δ*sarA* strains, each transformed with the pICS3 empty vector (/p), pICS3-P*amiA*-*rsaOI* (/p-*rsaOI*), or pICS3-P*amiA*-*rsaC* (/p-*rsaC*). Ready Blue staining of total protein extracts was used as loading control. For (B) and (C) cells were grown for 5 h in LB medium adjusted to pH 7 prior to protein extraction.

**Figure S6.**
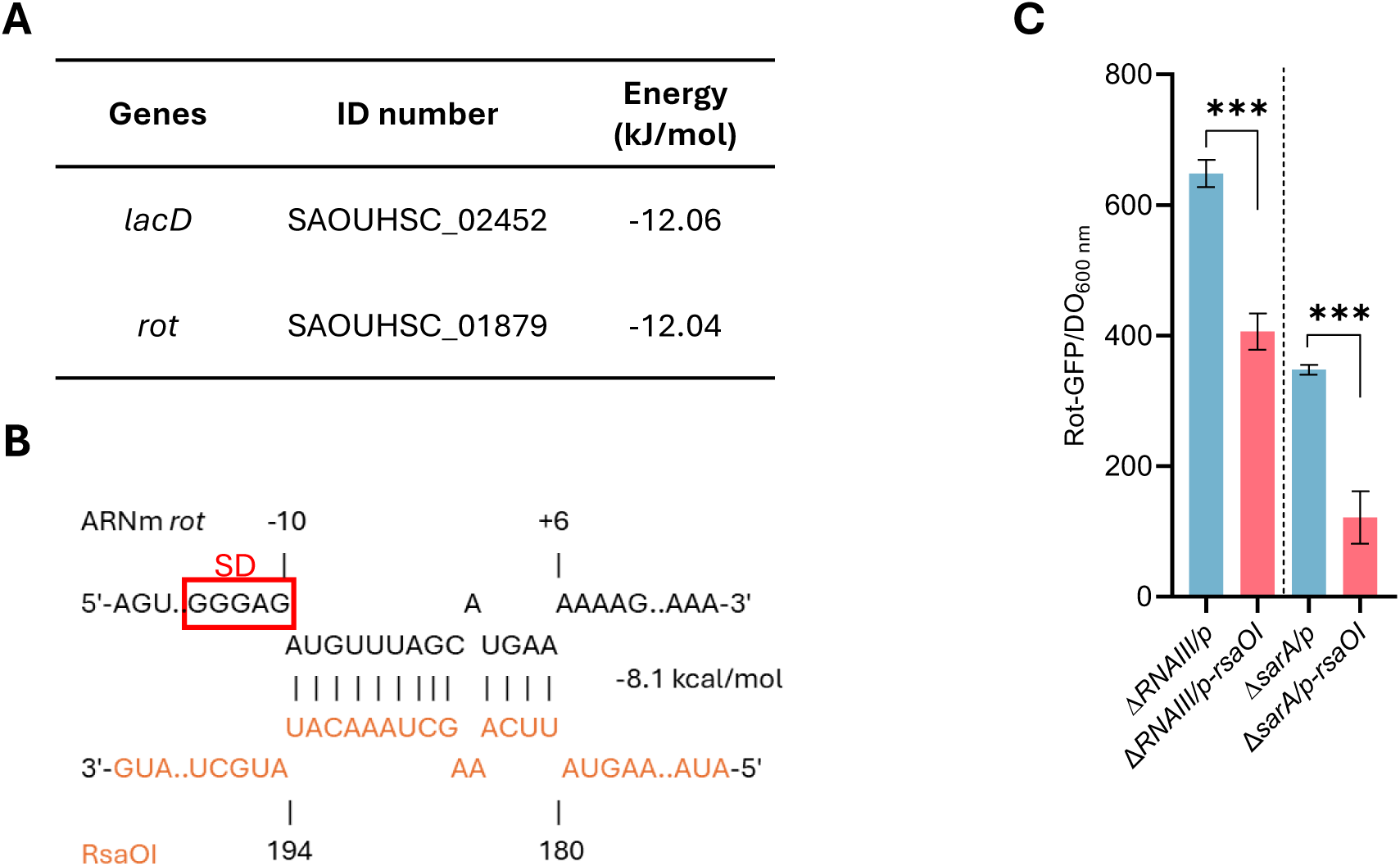
RsaOI modulate *rot* translation. (**A**) *In silico* RNA predator identify *rot* predicted RNA target of RsaOI immediately after *lacD* which is a known target of RsaOI. (**B**) IntaRNA prediction of interaction of RsaOI with *rot* mRNA, Shine-Dalgarno sequence is outlined in red. (**C**) The effect of RsaOI on *rot* expression was analyzed using *gfp* translational reporter assay, in HG003 Δ*RNAIII* and Δ*sarA* strains containing pICS3-empty vector (/*p*) and pICS3-*PamiA*-*rsaOI* (/*p-rsaOI*). The strains are co-transformed with pCN33-*PtufA*-*rot*-*gfp*. The fluorescence intensity was measured after 3h of growth in LB medium adjusted to pH7. Statistical analysis was conducted using Student’s *t*-test. Error bars represent the average of three independent experiments. Statistical significance is indicated by bars and asterisks as follows: \*\*\**P*< .001.

