## Supplementary material for "Cross-regulatory sRNA networks shape adaptive stress responses in *Staphylococcus aureus*: RsaC and RsaOI regulate galactose metabolism and cell wall homeostasis": Table S1, Table S2

**Table S1.** Strains and plasmids.

| **Strains** | **Relevant characteristics** | | **References** |
| --- | --- | --- | --- |
| ***E. coli* strains** | |  |  |
| DH5α | | *F⁻ φ80ΔlacZΔM15 Δ(lacZYA-argF)U169 recA1 endA1 hsdR17(rK⁻ mK⁺) phoA supE44 λ⁻ thi-1 gyrA96 relA1* | (1) |
| IM08b | | *SA08BΩPN25-hsdS (CC8-1)* (SAUSA300_0406) of NRS384 integrated between essQ and cspB genes | (2) |
| ***S. aureus* strains** | |  |  |
| RN4220 | | Restriction-defective derivative of 8325-4 | (3) |
| HG003 | | *rsbU* restored strain 8325-4, lysogenic for phages Φ11, Φ12, and Φ13 | (4) |
| HG003 Δ*vraR* | | *vraR* deleted HG003 (erythormycin resistance) | (5) |
| HG003 Δ*mntR* | | *mntR* deleted HG003 (erythormycin resistance) | This study |
| HG003 Δ*sarA* | | *sarA* deleted HG003 (kanamycin resistance) | This study |
| HG003 Δ*sarA*Δ*rot* | | *sarA* and *rot* deleted HG003 (kanamycin and erythromycin resistance) | This study |
| HG003 Δ*RNAIII* | | RNAIII deleted HG003 (kanamycin resistance) | (6) |
| HG003 Δ*sarS* | | *sarS* deleted HG003 (erythromycin resistance) | (7) |
| HG003 Δ*sarA*Δ*sarS* | | *sarA* and *sarS* deleted HG003 (kanamycin and erythromycin resistance) | This study |
| HG003 Δ*rsaOI* | | *rsaOI* deleted HG003 | (8) |
| HG003 Δ*srn_9342* | | *srn_9342* deleted HG003 | (9) |
| HG003 Δ*rsaC* | | *rsaC* deleted HG003 | (6) |
| HG003 Δ*rsaOI*ΔrsaC | | *rsaOI* and *rsaC* deleted HG003 | This study |
| SH1000 | | *rsbU* restored strain 8325-4 | (10) |
| SH1000 Δ*rot* | | *rot* deleted SH1000 | (10) |
| ***S. cerevisiae* strains** | |  |  |
| YM954 | | *MATa ade2 his3 leu2 lys2 trp1 ura3 gal4 gal80* | (11) |
| YAR_001 | | YM954 with the *sprC* promoter integrated into the *URA3* locus | This study |
| YAR_002 | | YM954 with the *srn_9340* promoter integrated into the *URA3* locus | This study |
| YAR_007 | | YM954 with the *srn_9342* promoter integrated into the *URA3* locus | This study |
| YAR_009 | | YM954 with the *sRNA71* promoter integrated into the *URA3* locus | This study |
| YAR_010 | | YM954 with the *rsaOI* promoter integrated into the *URA3* locus | This study |
| YAR_011 | | YM954 with the *rsaD* promoter integrated into the *URA3* locus | This study |
| YAR_012 | | YM954 with the *rsaC* promoter integrated into the *URA3* locus | This study |
| YAR_T | | YM954 without a promoter integrated at the *URA3* locus | This study |
| **Plasmids** | |  |  |
| pYI2267OZ | | Shuttle vector with *HIS3,* for integration of *psRNA-HIS3* into the *URA3 locus* | (12) |
| pYI2267OZ-*Psrn_9340* | | pYI2267OZ expressing *HIS3* under *srn_9340* promoter | This study |
| pYI2267OZ-*PsprC* | | pYI2267OZ expressing *HIS3* under *sprC* promoter | This study |
| pYI2267OZ-*Psrn_9342* | | pYI2267OZ expressing *HIS3* under *srn_9342* promoter | This study |
| pYI2267OZ-*PrsaD* | | pYI2267OZ expressing *HIS3* under *rsaD* promoter | This study |
| pYI2267OZ-*PsRNA71* | | pYI2267OZ expressing *HIS3* under *sRNA71* promoter | This study |
| pYI2267OZ-*PrsaOI* | | pYI2267OZ expressing *HIS3* under *rsaOI* promoter | This study |
| pYI2267OZ-*PrsaC* | | pYI2267OZ expressing *HIS3* under *rsaC* promoter | This study |
| pCN41c | | Shuttle vector with *blaZ* | (7) |
| pCN41c-*Psrn_9340* | | pCN41c expressing *blaZ* under *srn_9340* promoter | This study |
| pCN41c-*PsprC* | | pCN41c expressing *blaZ* under *sprC* promoter | (7) |
| pCN41c-*PsRNA71* | | pCN41c expressing *blaZ* under *sRNA71* promoter | This study |
| pCN41c-*Psrn_9342* | | pCN41c expressing *blaZ* under *srn_9342* promoter | This study |
| pCN41c-*PrsaD* | | pCN41c expressing *blaZ* under *rsaD* promoter | This study |
| pICS3 | | Shuttle vector, Cm^r^ (*cat194*), pC194 replicon | (13) |
| pICS3-*PamiA-rsaOI* | | pICS3 with *rsaOI* of HG003 under control of a consitutive promoter *amiA* | (5) |
| pICS3-*PamiA-rsaC* | | pICS3 with *rsaC* of HG003 under control of a consitutive promoter *amiA* | This study |
| pCN51-*P3* | | Shuttle vector with P3 promoter | (14) |
| pCN51-*P3*-*rsaC* | | pCN51 expressing *rsaC* under P3 promoter | (15) |
| pCN33-*PtufA*-*rot*-*gfp* | | pCN33 with *rot* of HG003 fused to *gfp* under control of *PtufA* promoter | (16) |
| pACTII | | Shuttle vector with *GAL4 AD* from *S. cereveisiae* | (17) |
| pACTII-*sarA* | | pACTII with *sarA* translationally fused to the *GAL4 AD* under the control of the constitutive promoter *PADH1* | This study |
| pACTII-c*odY* | | pACTII with *codY* translationally fused to the *GAL4 AD* under the control of the constitutive promoter *PADH1* | This study |
| pACTII-s*arS* | | pACTII with *sarS* translationally fused to the *GAL4 AD* under the control of the constitutive promoter *PADH1* | This study |
| pACTII-*purR* | | pACTII with *purR* translationally fused to the *GAL4 AD* under the control of the constitutive promoter *PADH1* | This study |
| pACTII-*mgrA* | | pACTII with *mgrA* translationally fused to the *GAL4 AD* under the control of the constitutive promoter *PADH1* | This study |
| pACTII-*ccpA* | | pACTII with *ccpA* translationally fused to the *GAL4 AD* under the control of the constitutive promoter *PADH1* | This study |
| pACTII-*agrA* | | pACTII with *agrA* translationally fused to the *GAL4 AD* under the control of the constitutive promoter *PADH1* | This study |
| pACTII-*walR* | | pACTII with *walR* translationally fused to the *GAL4 AD* under the control of the constitutive promoter *PADH1* | This study |
| pACTII-*mntR* | | pACTII with *mntR* translationally fused to the *GAL4 AD* under the control of the constitutive promoter *PADH1* | This study |
| pACTII-*rot* | | pACTII with *rot* translationally fused to the *GAL4 AD* under the control of the constitutive promoter *PADH1* | This study |
| pACTII-*sarU* | | pACTII with *sarU* translationally fused to the *GAL4 AD* under the control of the constitutive promoter *PADH1* | This study |
| pACTII-*icaR* | | pACTII with *icaR* translationally fused to the *GAL4 AD* under the control of the constitutive promoter *PADH1* | This study |
| pACTII-*fur* | | pACTII with *fur* translationally fused to the *GAL4 AD* under the control of the constitutive promoter *PADH1* | This study |
| pACTII-*sarV* | | pACTII with *sarV* translationally fused to the *GAL4 AD* under the control of the constitutive promoter *PADH1* | This study |
| pACTII-*sarR* | | pACTII with *sarR* translationally fused to the *GAL4 AD* under the control of the constitutive promoter *PADH1* | This study |
| pACTII-*sarT* | | pACTII with *sarT* translationally fused to the *GAL4 AD* under the control of the constitutive promoter *PADH1* | This study |
| pACTII-*pyrR* | | pACTII with *pyrR* translationally fused to the *GAL4 AD* under the control of the constitutive promoter *PADH1* | This study |
| pACTII-*fapR* | | pACTII with *fapR* translationally fused to the *GAL4 AD* under the control of the constitutive promoter *PADH1* | This study |
| pACTII-*phoU* | | pACTII with *phoU* translationally fused to the *GAL4 AD* under the control of the constitutive promoter *PADH1* | This study |
| pACTII-*arlR* | | pACTII with *arlR* translationally fused to the *GAL4 AD* under the control of the constitutive promoter *PADH1* | This study |
| pACTII-*graR* | | pACTII with *graR* translationally fused to the *GAL4 AD* under the control of the constitutive promoter *PADH1* | This study |
| pACTII-*srrA* | | pACTII with *srrA* translationally fused to the *GAL4 AD* under the control of the constitutive promoter *PADH1* | This study |
| pACTII-*cymR* | | pACTII with *cymR* translationally fused to the *GAL4 AD* under the control of the constitutive promoter *PADH1* | This study |
| pACTII-*nrdR* | | pACTII with *nrdR* translationally fused to the *GAL4 AD* under the control of the constitutive promoter *PADH1* | This study |
| pACTII-*perR* | | pACTII with *perR* translationally fused to the *GAL4 AD* under the control of the constitutive promoter *PADH1* | This study |
| pACTII-*vraR* | | pACTII with *vraR* translationally fused to the *GAL4 AD* under the control of the constitutive promoter *PADH1* | This study |
| pACTII-*pitR* | | pACTII with *pitR* translationally fused to the *GAL4 AD* under the control of the constitutive promoter *PADH1* | This study |
| pACTII-*cidR* | | pACTII with *cidR* translationally fused to the *GAL4 AD* under the control of the constitutive promoter *PADH1* | This study |
| pACTII-*gbaA* | | pACTII with *gbaA* translationally fused to the *GAL4 AD* under the control of the constitutive promoter *PADH1* | This study |
| pACTII-*argR* | | pACTII with *argR* translationally fused to the *GAL4 AD* under the control of the constitutive promoter *PADH1* | This study |
| pACTII-*arcR* | | pACTII with *arcR* translationally fused to the *GAL4 AD* under the control of the constitutive promoter *PADH1* | This study |
| pACTII-*hssR* | | pACTII with *hssR* translationally fused to the *GAL4 AD* under the control of the constitutive promoter *PADH1* | This study |
| pACTII-*rsbV* | | pACTII with *rsbV* translationally fused to the *GAL4 AD* under the control of the constitutive promoter *PADH1* | This study |
| pACTII-*lytR* | | pACTII with *lytR* translationally fused to the *GAL4 AD* under the control of the constitutive promoter *PADH1* | This study |
| pACTII-*spxA* | | pACTII with *spxA* translationally fused to the *GAL4 AD* under the control of the constitutive promoter *PADH1* | This study |
| pACTII-*mraZ* | | pACTII with *mraZ* translationally fused to the *GAL4 AD* under the control of the constitutive promoter *PADH1* | This study |
| pACTII-*hptR* | | pACTII with *hptR* translationally fused to the *GAL4 AD* under the control of the constitutive promoter *PADH1* | This study |
| pACTII-*lrgA* | | pACTII with *lrgA* translationally fused to the *GAL4 AD* under the control of the constitutive promoter *PADH1* | This study |
| pACTII-*yabJ* | | pACTII with *yabJ* translationally fused to the *GAL4 AD* under the control of the constitutive promoter *PADH1* | This study |
| pACTII-*spoVG* | | pACTII with *spoVG* translationally fused to the *GAL4 AD* under the control of the constitutive promoter *PADH1* | This study |
| pACTII-*saeR* | | pACTII with *saeR* translationally fused to the *GAL4 AD* under the control of the constitutive promoter *PADH1* | This study |
| pACTII-*comK1* | | pACTII with *comK1* translationally fused to the *GAL4 AD* under the control of the constitutive promoter *PADH1* | This study |
| pACTII-*zur* | | pACTII with *zur* translationally fused to the *GAL4 AD* under the control of the constitutive promoter *PADH1* | This study |

**Table S2.** Oligonucleotides used in this study.

| **Oligonucleotides** | **Sequences** | **Experiment** |
| --- | --- | --- |
| T7_*rsaOI*_5’ | TAATACGACTCACTATAGGGATAGAGAAAAGAAGTTATCGC | EMSA |
| T7_*rsaOI*_3’ | AACCCCCACTGCAATGATTATCG | EMSA |
| T7_*rsaC*_544_5’ | TAATACGACTCACTATAGGGAAAAGCTTTATGTGG | EMSA |
| T7_*rsaC*_544_3’ | AAAATAGCCACACTCATATG | EMSA |
| T7_*rsaC*_1116_5’ | TAATACGACTCACTATAGGGCATATCGTGCGTTAAATATATATTAC | EMSA |
| T7_*rsaC*_1116_3’ | AAAATAGCCACACTCATATG | EMSA |
| T7_*srn_9342*_short_5’ | GCCAGTTAATACGACTCACTATAGGGGGAGGGGCAACGTTATTA | EMSA |
| T7_*srn_934*2_short_3’ | GTAAATAGAAAGCAGGTATGTAAC | EMSA |
| T7_*srn_9342*_long_5’ | GCCAGTTAATACGACTCACTATAGGGGGAGGGGCAACGTTATTA | EMSA |
| T7_*srn_9342*_long_3’ | AAAAATAGGCAAGTACCGTAG | EMSA |
| T7_*lacC*_5’ | TAATACGACTCACTATAGGGCTAAAATTAAACATTTA | EMSA |
| T7_*lacC*_3’ | CCTTTACCACCAGCTGTT | EMSA |
| Srn_9342 | GTAACGCACCTGCTTAAATAGACAT | Northern Blot |
| RsaOI | TGAGACTCTAACCTTCAATAT | Northern Blot |
| RsaC | GGGCATACAATTGAACAATCTGTAATAG | Northern Blot |
| Srn_9340 | CGACTAACTCCTGGTTATGC | Northern Blot |
| SprC | CATATATATATTTACTATGAAC | Northern Blot |
| sRNA71 | GGATGACGTTGGGTTGCGCCATCC | Northern Blot |
| ARNtm | ACACGCTTAATGAGCTCGGG | Northern Blot |
| qPCR_*gyrB*_5’ | TGAAGTGCGTCAAGTTGTAG | RT-qPCR |
| qPCR_*gyrB*_ 3’ | TGCGACTTGTGGATTTTCAT | RT-qPCR |
| qPCR_ *sarA*_5’ | TTCGTTGTTTGCTTCAGTGA | RT-qPCR |
| qPCR _*sarA*_3’ | AAAAGAAGCGCGCGATAAAG | RT-qPCR |
| qPCR_*tagO*_5’ | TGTACGTACTTGGGCTTGTAGA | RT-qPCR |
| qPCR_*tagO*_3’ | AAGCAACTACTAAGGCAGCG | RT-qPCR |
| qPCR _*murD* _5’ | GCAACAAACACGCTAGCAAC | RT-qPCR |
| qPCR_*murD*_3’ | TCGCGCGAACATTTTCCATA | RT-qPCR |
| qPCR_*murG*_5’ | GTAGAGCTGGATCAAATGCGA | RT-qPCR |
| qPCR_*murG*_3’ | GTCGCCTCGGGATTGATCTA | RT-qPCR |
| qPCR_*02820*_5’ | CTCTAGTTTCCAAAGCGGGA | RT-qPCR |
| qPCR_*02820*_3’ | GCTGAAACGCGGACAAATAG | RT-qPCR |
| qPCR _*sscB*_5’ | AGTTTTATTGGACCACGCCC | RT-qPCR |
| qPCR_*sscB*_3’ | TCCACCTTGAATTTGCGCAA | RT-qPCR |
| qPCR_*rot*_5’ | GCGTCCTGTTGACGATGAAA | RT-qPCR |
| qPCR_*rot*_3’ | TGCATTGCTGTTGCTCTACTTG | RT-qPCR |
| qPCR _*dltA*_5’ | GTTTCCCAAGTGCGACGATT | RT-qPCR |
| qPCR _*dltA*_3’ | TGGTCTTTCAACGCCAACAG | RT-qPCR |
| qPCR _*pbp2*_5’ | TTTCTACGAACATGGCGCAC | RT-qPCR |
| qPCR _ *pbp2*_3’ | TGAGGCACCTTCAGAACCAA | RT-qPCR |
| qPCR_*mprF*_5’ | TCGAACGTTGGTCAATTGCA | RT-qPCR |
| qPCR_*mprF*_3’ | TTCAAAGACACGGCCTGCAA | RT-qPCR |
| qPCR _*tarL*_5’ | ACTTGGGAAGGCAGACTACA | RT-qPCR |
| qPCR_*tarL*_3’ | TTTTAAAGGCACCAACGGCA | RT-qPCR |
| qPCR_*sarS*_5’ | CAAGAAAACACACTTCCATT | RT-qPCR |
| qPCR_*sarS*_3’ | TATATGCTGTACTAAATCCG | RT-qPCR |
| qPCR_*spa*_5’ | CGCAACACGATGAAGCTC | RT-qPCR |
| qPCR_*spa*_3’ | ACCATTGCGTTGATCAGCAT | RT-qPCR |
| qPCR_*agrA*_5' | CGAAGACGATCCAAAACAAAGAG | RT-qPCR |
| qPCR_*agrA*_3' | GCTCAAGCACCTCATAAGGAT | RT-qPCR |
| qPCR_*psmα*_5' | TATCATCGCTGGCATCATTA | RT-qPCR |
| qPCR_*psmα*_3' | AGACCTCCTTTGTTTGTTATG | RT-qPCR |
| qPCR_*rsaC*_5' | GCCATTCCCTACACACTCTTAT | RT-qPCR |
| qPCR_*rsaC*_3' | CAGTTGTCCAATCACATAAGAACC | RT-qPCR |
| *Psrn_9340_*XhoI_5' | CGCGCTCGAGCAAGATTTTGAAGTTTAGT | Cloning |
| *Psrn_9340_*XhoI_3' | CGCGCTCGAGGATTATATTATAATATTATT | Cloning |
| *Psrn_9342_*XhoI_5' | CGCGCTCGAGAAGTGCATGGAAACGTAATAAATATG | Cloning |
| *Psrn_9342_*XhoI_3' | CGCGCTCGAGCCCCTCCCATGTATATCCTAC | Cloning |
| *PsRNA71_*XhoI_5' | CCGGCTCGAGACTTTGACTACTAATTCG | Cloning |
| *PsRNA71_*XhoI_3' | GGCCCTCGAGGACTGTGTAGTCATTTAAG | Cloning |
| *PrsaOI_*XhoI_5' | CCGGCTCGAGATATGTAATGCTATGTCATTAG | Cloning |
| *PrsaOI_*XhoI_3' | GGCCCTCGAGATACATATAGTAACGTCTTATC | Cloning |
| *PrsaC_*XhoI_5' | CCGGCTCGAGTTAACATACTTTCACCTCAC | Cloning |
| *PrsaC_*XhoI_3' | GGCCCTCGAGTACCTCCTAATTAAAAGTTT | Cloning |
| *PrsaD_*XhoI_5' | CGCGCTCGAGCCTTTATATATTTATTAGAC | Cloning |
| *PrsaD_*XhoI_3' | CGCGCTCGAGGAAGTTAAAAACATTATAGC | Cloning |
| *PsRNA71*_BamHI_5' | CCGGGGATCCACTTTGACTACTAATTCG | Cloning |
| *PsRNA71*_EcoRI_3' | GGCCGAATTCGACTGTGTAGTCATTTAAG | Cloning |
| *PrsaD*_BAMHI_5' | GGCCGGATCCCACGAAACCATCCCCTTTA | Cloning |
| *PrsaD*_EcoRI_3' | GGCCGAATTCGTTAAAAACATTATAGCAGACA | Cloning |
| *Psrn9340*_BamHI_5' | GGCCGGATCCATCAAATGTGAATATGTTTTAAGCG | Cloning |
| *Psrn9340*_EcoRI_3' | GCGCGAATTCGATTATATTATAATATTATTA | Cloning |
| *Psrn_9342*_BamHI_5' | TCAGGGATCCCTAGATTCACAAACGGCAATCAACC | Cloning |
| *Psrn_9342*_EcoRI_3' | TTCAGGAATTCTAGGCAAGTAATAACGTTGC | Cloning |
| pICS3-*PamiA*-*rsaC*-5' | TTAAAAGCTTGCATGCCTGCAGGAAAATTTGTTTGATTTTTAATGGATAATGTGATATAATGGCATATCGTGCGTTAAATATATATTAC | Cloning |
| pICS3-*PamiA*-*rsaC*-3' | GAGAAAATACCGCATCAGGCGCCGGTGAAGTTAAGTGTATTCAC | Cloning |
| NcoI_*sarA_*5' | CCGGGCCATGGCAATTACAAAAATCAATG | Cloning |
| BamHI_*sarA_*3 | GGCCGGATCCTTATAGTTCAATTTCGTTGTTTGC | Cloning |
| NcoI_*codY_*5' | CCGGCCATGGGAATGAGCTTATTATCTAAAAC | Cloning |
| BamHI_*codY_*3' | GGCCGGATCCTTATTTACTTTTTTCTAATTC | Cloning |
| NcoI_*sarS_*5' | CCGGGCCATGGGAATGAAATATAATAACCATGAC | Cloning |
| BamHI_*sarS*_3' | GGCCGGATCCTTATTCAAAAACAAGATGTAAATG | Cloning |
| NcoI_*purR*_5' | CCGGGCCATGGGAATGAGATATAAACGAAGCGAG | Cloning |
| BamHI_*purR*_3' | GGCCGGATCCTTATGAAAACTTAGATAAACTGTTG | Cloning |
| NcoI_*mgrA*_5' | CCGGGCCATGGGAATGTCTGATCAACATAATTTAAAAG | Cloning |
| BamHI_m*grA*_3' | GGCCGGATCCTTATTTTTCCTTTGTTTCATCAAATGC | Cloning |
| NcoI_*ccpA_5'* | CCGGGCCATGGGAATGACAGTTACTATATATGATG | Cloning |
| BamHI_*ccpA*_3' | GGCCGGATCCTTATTTTGTAGTTCCTCGGTATTC | Cloning |
| NcoI_*agrA*_5' | CCGGGCCATGGAAATTGCCCTCGCAAC | Cloning |
| BamHI_*agrA*_3' | GGCCGGATCCTTATATTTTTTTAACGTTTCTCACCG | Cloning |
| NcoI_*walR*_5' | CCGGGCCATGGGAATGGCTAGAAAAGTTGTTGTAG | Cloning |
| BamHI_*walR*_3' | GGCCGGATCCCTACTCATGTTGTTGGAGGAAATAT | Cloning |
| NcoI_*mntR*_5' | CCGGGCCATGGGAATGTTAACTGAAGAAAAAGAGG | Cloning |
| BamHI_*mntR*_3' | GGCCGGATCCTTATTTTTCAGCAAACATGTTCATTG | Cloning |
| NcoI_*rot*_5' | CCGGGCCATGGGAATGAAAAAAGTAAATAACGACACT | Cloning |
| BamHI_*rot*_3' | GGCCGGATCCTTACACAGCAATAATTGCGTTTAAAC | Cloning |
| NcoI_*sarU*_5' | CCGGGCCATGGGAGTGGACTATCAAACTTTCGAAA | Cloning |
| BamHI_*sarU*_3' | GGCCGGATCCTTAAAAGAAAAATTTTCTTGGTTTGAG | Cloning |
| NcoI_*icaR*_5' | CCGGGCCATGGGATTGAAGGATAAGATTATTGATAAC | Cloning |
| BamHI_*icaR*_3' | GGCCGGATCCTTATTTCTTCAAAAATATATTTAGTAGC | Cloning |
| EcoRI_*fur*_5' | CCGGGGAATTCGATTGGAAGAACGATTAAATCGC | Cloning |
| XhoI_*fur*_3' | GGCCCTCGAGCTATCCTTTACCTTTAGCTTGGCA | Cloning |
| NcoI_s*arV*_5' | CCGGGCCATGGGAATGAGTAATAAAGTTCAACGTTTT | Cloning |
| BamHI_*sarV*_3' | GGCCGGATCCTTAGTTTGTAACGCTTAACACTTCT | Cloning |
| NcoI_*sarR*_5' | CCGGGCCATGGGAATGAGTAAAATTAATGACATTAATG | Cloning |
| BamHI_*sarR*_3' | GGCCGGATCCTTAATTTTTAATGTATTCTTCTAATTCT | Cloning |
| NcoI_*sarT*_5' | CCGGGCCATGGGAATGAAAAGAGTATTAACTACATAT | Cloning |
| BamHI_*sarT*_3' | GGCCGGATCCTTATGTTTCATTAATATTTATTTCACTC | Cloning |
| NcoI_*pyrR*_5' | CCGGCCATGGGAATGTCTGAACGTATCATAATG | Cloning |
| BamHI_*pyrR*_3' | GGCCGGATCCTTATTTAATTATAACTGCATTTC | Cloning |
| NcoI_*fapR*_5' | CCGGCCATGGGAATGAGGGGTGAGACGTTGAAAC | Cloning |
| BamHI_*fapR*_3' | GGCCGGATCC TTATCCTCGCTTATCATAAAAC | Cloning |
| NcoI_*phoU*_5' | CCGGCCATGGGAATGGCAATAATTAGACAACG | Cloning |
| BamHI_*phoU*_3' | GGCCGGATCCTTATTGTTCGTAATGTGTAC | Cloning |
| NcoI_*arlR*_5' | CCGGCCATGGGAATGACGCAAATTTTAATAG | Cloning |
| BamHI_*arlR*_3' | GGCCGGATCCTCATCGTATCACATACCC | Cloning |
| NcoI_*graR*_5' | CCGGCCATGGGAATGCAAATACTACTAGTAGAAG | Cloning |
| BamHI_*graR*_3' | GGCCGGATCCTTATTCATGAGCCATATATCC | Cloning |
| NcoI_*srrA*_5' | CCGGCCATGGGAATGTCGAACGAAATACTTATC | Cloning |
| BamHI_*srrA*_3' | GGCCGGATCCCTATTTAGCCGGCTCATCATTAG | Cloning |
| NcoI_*cymR*_5' | CCGGCCATGGGAATGAAAATTTCTACTAAAG | Cloning |
| BamHI_*cymR*_3' | GGCCGGATCCTTAAATATAAAACATGTATC | Cloning |
| NcoI_*nrdR*_5' | CCGGCCATGGGAATGAAATGCCCGAAATG | Cloning |
| BamHI_*nrdR*_3' | GGCCGGATCCTTAAGCATCACTCCGTTTG | Cloning |
| NcoI_*perR*_5' | CCGGCCATGGGAATGAGTGTTGAAATAGAATC | Cloning |
| BamHI_*perR*_3' | GGCCGGATCCTTATTTATCTTGGCATTCTTTAC | Cloning |
| NcoI_v*raR*_5' | CCGGCCATGGGAATGACGATTAAAGTATTG | Cloning |
| BamHI_*vraR*_3' | GGCCGGATCCCTATTGAATTAAATTATG | Cloning |
| NcoI_*pitR*_5' | CCGGCCATGGGAATGTTTAGTAAGAAAAAAG | Cloning |
| BamHI_*pitR*_3' | GGCCGGATCCTTAGCTATTTTTCATAATAATAG | Cloning |
| NcoI_*cidR*_5' | CCGGCCATGGGAATGACTAATGCTTCCAAATC | Cloning |
| BamHI_*cidR*_3' | GGCCGGATCCTTAGCCTAAACGGTCTTTC | Cloning |
| NcoI_*gbaA*_5' | CCGGCCATGGGAATGCGAAAAGATGCAAAAG | Cloning |
| BamHI_*gbaA*_3' | GGCCGGATCCTTAGTCATTACGTCCCACC | Cloning |
| NcoI_*argR*_5' | CCGGCCATGGGAGTGCCCAAAAAATCGGTTAG | Cloning |
| BamHI_*argR*_3' | GGCCGGATCCTTATAACAAATTGAAAATTC | Cloning |
| NcoI_*arcR*_5' | CCGGCCATGGGAATGACAGAAAACTTTATTTTG | Cloning |
| BamHI_*arcR*_3' | GGCCGGATCCTTAAACACATACATCATTG | Cloning |
| NcoI_*hssR*_5' | CCGGCCATGGGAATGGTGCAATGTCTTGTTG | Cloning |
| BamHI_*hssR*_3' | GGCCGGATCCTTAAACATGATTCTCCACC | Cloning |
| NcoI_*rsbV*_5' | CCGGCCATGGGAATGAATCTTAATATAGAAAC | Cloning |
| BamHI_*rsbV*_3' | GGCCGGATCCTTATTCGACCTCCGTTCCTTC | Cloning |
| NcoI_*lytR*_5' | CCGGCCATGGGAATGAAAGCATTAATCATAG | Cloning |
| BamHI_*lytR*_3' | GGCCGGATCCTTAAAGTAATCCTATCGAC | Cloning |
| NcoI_*spxA*_5' | CCGGCCATGGGAATGGTAACATTATTTACTTC | Cloning |
| BamHI_*spxA*_3' | GGCCGGATCCTTAGTCAACCATACGTTGTG | Cloning |
| NcoI_*mraZ*_5' | CCGGCCATGGGAATGTTCATGGGAGAATACGATC | Cloning |
| BamHI_*mraZ*_3' | GGCCGGATCCTTAAAAATCAAAATCTATTAAATC | Cloning |
| NcoI_*hptR*_5' | CCGGCCATGGGAATGTTTAAGGTAGTTATTTGTG | Cloning |
| BamHI_*hptR*_3' | GGCCGGATCCCTATTTTGCTTGCTTACAATAATC | Cloning |
| NcoI_*lrgA*_5' | CCGGCCATGGGAATGGTCGTGAAACAACAAAAAG | Cloning |
| BamHI_*lrgA*_3' | GGCCGGATCCTTAATCATGAGCTTGTGCCTCC | Cloning |
| NcoI_*yabJ*_5' | CCGGCCATGGGAATGAAAATCATTAACACAAC | Cloning |
| BamHI_*yabJ*_3' | GGCCGGATCCTTATAATTCCTTAATTTTAC | Cloning |
| NcoI_*spoVG*_5' | CCGGCCATGGGAATGAAAGTGACAGATGTAAG | Cloning |
| BamHI_*spoVG*_3' | GGCCGGATCCTTAAGCTTCTTCTGAATCTTC | Cloning |
| NcoI_*saeR*_5' | CCGGCCATGGGAATGACCCACTTACTGATCGTG | Cloning |
| BamHI_*saeR*_3' | GGCCGGATCCTTATCGGCTCCTTTCAAATTT | Cloning |
| NcoI_*comK1*_5' | CCGGCCATGGGAATGGTTATTCGACCAGCATTTG | Cloning |
| BamHI_*comK1*_3' | GGCCGGATCCTTATTTATCTTCAAATAAAG | Cloning |
| NcoI_*zur*_5' | CCGGCCATGGGAATGAATACAAATGATGC | Cloning |
| BamHI_*zur*_3' | GGCCGGATCCTTAATCTTGGCAAGACTC | Cloning |
